# Stoichiometric expression of messenger polycistrons by eukaryotic ribosomes (SEMPER) for compact, ratio-tunable multi-gene expression from single mRNAs

**DOI:** 10.1101/2023.05.26.541240

**Authors:** Mengtong Duan, Ishaan Dev, Andrew Lu, Mei Yi You, Mikhail G. Shapiro

## Abstract

Applications of mammalian synthetic biology increasingly require the ability to express multiple proteins at user-determined stoichiometries from single, compactly encoded transcripts. Here we present an approach for expressing multiple open reading frames (ORFs) from a single transcript, taking advantage of the leaky scanning model of translation initiation. In this method, adjacent ORFs are translated from a single messenger RNA at tunable ratios determined by their order in the sequence and the strength of their translation initiation sites. We call this approach Stoichiometric Expression of Messenger Polycistrons by Eukaryotic Ribosomes (SEMPER). We demonstrate the principles of this approach by expressing up to three fluorescent proteins from one plasmid in two different cell lines. We then use it to encode a stoichiometrically tuned polycistronic construct encoding gas vesicle acoustic reporter genes, showing that enforcing the optimal ratio in every cell enables efficient formation of the multi-protein complex while minimizing cellular toxicity. Finally, we demonstrate the polycistronic expression of two fluorescent proteins from single mRNAs made through *in vitro* transcription and delivered to cells. SEMPER will enable a broad range of applications requiring tunable expression from compact eukaryotic constructs.

## INTRODUCTION

Mammalian cell engineering promises to enable treatment of age-related degeneration, reversal of genetic diseases, and even transformation of cells into living therapeutics and sensors.^1^ Realizing this promise requires the development of genetic circuits capable of finely tuning the relative expression stoichiometries of multiple proteins to produce functional multimeric protein assemblies, multi-component signaling systems, or multi-enzyme biosynthetic pathways.^2–6^ Current approaches to doing so at the DNA level (e.g., varying promoter strength or titrating copy numbers of each gene) yield lengthy DNA constructs that often must be packaged into multiple delivery vectors and lead to undesirable cell-to-cell variability due to stochastic gene delivery and integration across the population. Additionally, attainable protein expression stoichiometries are limited by the transcriptional strengths of a relatively small set of curated promoters that often demonstrate cell-to-cell variability.^7^

Post-transcriptionally, sequence motifs such as internal ribosome entry sites (IRES) or 2A self-cleaving peptides may be used to encode multiple open reading frames (ORF) into a single transcript and tune protein stoichiometries using relative translation levels.^8,9^ Such post-transcriptional mechanisms are also useful for mRNA-based, multi-gene expression systems, which are of relevance to mRNA vaccine and therapeutic development.^10^ Although powerful tools, these genetic parts have significant disadvantages. For instance, IRES sequences have a significant genetic footprint (∼200-600 bps),^11^ leading to lengthier genetic constructs which may reduce viral packaging efficiency. Likewise, 2A self-cleaving peptides leave peptide scars and can yield undesired fusion proteins, both of which can be detrimental to protein function.^12^ In addition, when employed in bicistronic vectors, these genetic parts can strongly attenuate the expression of the second ORF relative to the expression of the first ORF.^9,13^ Here, we present an alternative approach that enables compact and robustly tunable polycistronic expression in mammalian cells using plasmid- and mRNA-based expression systems. We call this approach “Stoichiometric Expression of Messenger Polycistrons by Eukaryotic Ribosomes” (SEMPER).

SEMPER uses the canonical cap-dependent ribosome recruitment and translation mechanism in mammalian systems, which begins when the 43S preinitiation complex (PIC) of the ribosome is loaded onto the 5’ end of mRNA.^14,15^ This complex then scans the 5’ untranslated region (5’UTR) until it encounters a translation initiation site (TIS), consisting of the start codon (AUG) and ∼3-10 neighboring nucleotides.^16^ The TIS sets the translational reading frame and initiates translation by engaging with the 60S ribosomal subunit. The full ribosome then translates the mRNA into protein until it encounters a stop codon, where it terminates translation and disengages the transcript. With some frequency, the 43S PIC may scan through the first TIS and initiate translation from a downstream ORF starting at another TIS, a phenomenon called leaky ribosomal scanning (LRS).^17^ As determined by its sequence, a strong TIS (e.g. the Kozak consensus sequence) will reliably initiate translation while weaker ones will more frequently allow the 43S PIC to scan past.^18^

By employing a short ORF (uORF) upstream of a gene of interest (GOI), mammalian cells naturally use LRS and alternate TISs to divert a portion of the ribosome flux away from a GOI, effectively downregulating its translation.^19^ Recently, Ferreira and colleagues demonstrated that it is possible to use synthetic uORFs to regulate the expression of a downstream recombinant GOI.^20^ They also empirically determined the strength of various translation initiation sequences and showed that it is possible to divert varying amounts of ribosomal flux away from the GOI by varying the uORF TIS strength. The SEMPER approach replaces non-protein-coding uORFs with longer, functional coding sequences. In this study, we show that by chaining together multiple ORFs while varying the translation initiation strength of each ORF, this approach achieves polycistronic expression of multiple proteins with tunable translation rates. We further demonstrate that this framework is functional in *in vitro* transcribed mRNA, paving the way for advances in mRNA-based protein therapeutics of higher complexity.

## RESULTS

### Tunable, plasmid-based SEMPER framework for expressing two ORFs

To test tunable translation levels of two recombinant proteins from single transcripts, we encoded fluorescent proteins (FPs) with minimal spectral overlap into the first two ORFs of our SEMPER plasmid vector (**Figure 1A**). We included 11 bps of distance between these ORFs to ensure that read-through of the stop codon of the first ORF would not result in a fusion containing both proteins. We used the following TIS sequence (NNNAUGG) to initiate translation of our ORFs, where NNN represents one of the following sequences in order of decreasing translation initiation efficiency: ACC, CCC, TTT.^20^ We engineered all our FPs to contain a valine residue (GTG) following the N-terminal methionine to ensure changes in translation initiation were due to the trinucleotide preceding the start codon. TIS sequences will be referred to by their variable NNN sequence. We used one other sequence (TTTCCAT), referred to as ***, to scrub the TIS entirely and prevent the ORF from being translated. For the first ORF, we generated a methionine-less monomeric Superfolder GFP (msfGFP[r5M]) in which all methionines—except the N-terminal one— were mutated to other amino acids.^21,22^ In addition, out-of-frame AUGs were removed from the msfGFP[r5M] coding sequence using synonymous mutations. These mutations effectively removed all internal TISs within ORF 1 that could reduce ribosomal flux to downstream ORFs (**Figure 1B**). In the second ORF, we encoded mEBFP2 with all its natural methionines. Finally, downstream of the SEMPER ORFs, we included an IRES followed by mCherry (IRES-mCherry) for normalization. As ribosomal binding to the IRES and subsequent translation of mCherry are conducted independently of ORF 1 and ORF 2 translation,^23,24^ the mCherry allowed us to normalize single-cell fluorescence measurements for msfGFP[r5M] and mEBFP2, a strategy common to LRS-focused studies.^20,25^ Because the analyte ORFs and IRES-mCherry were encoded on the same transcript, this normalization scheme accounted for variations in transfection efficiency, transcription, and mRNA decay. mCherry fluorescence also served as a proxy for transcript abundance in each cell.

**Figure 1:**
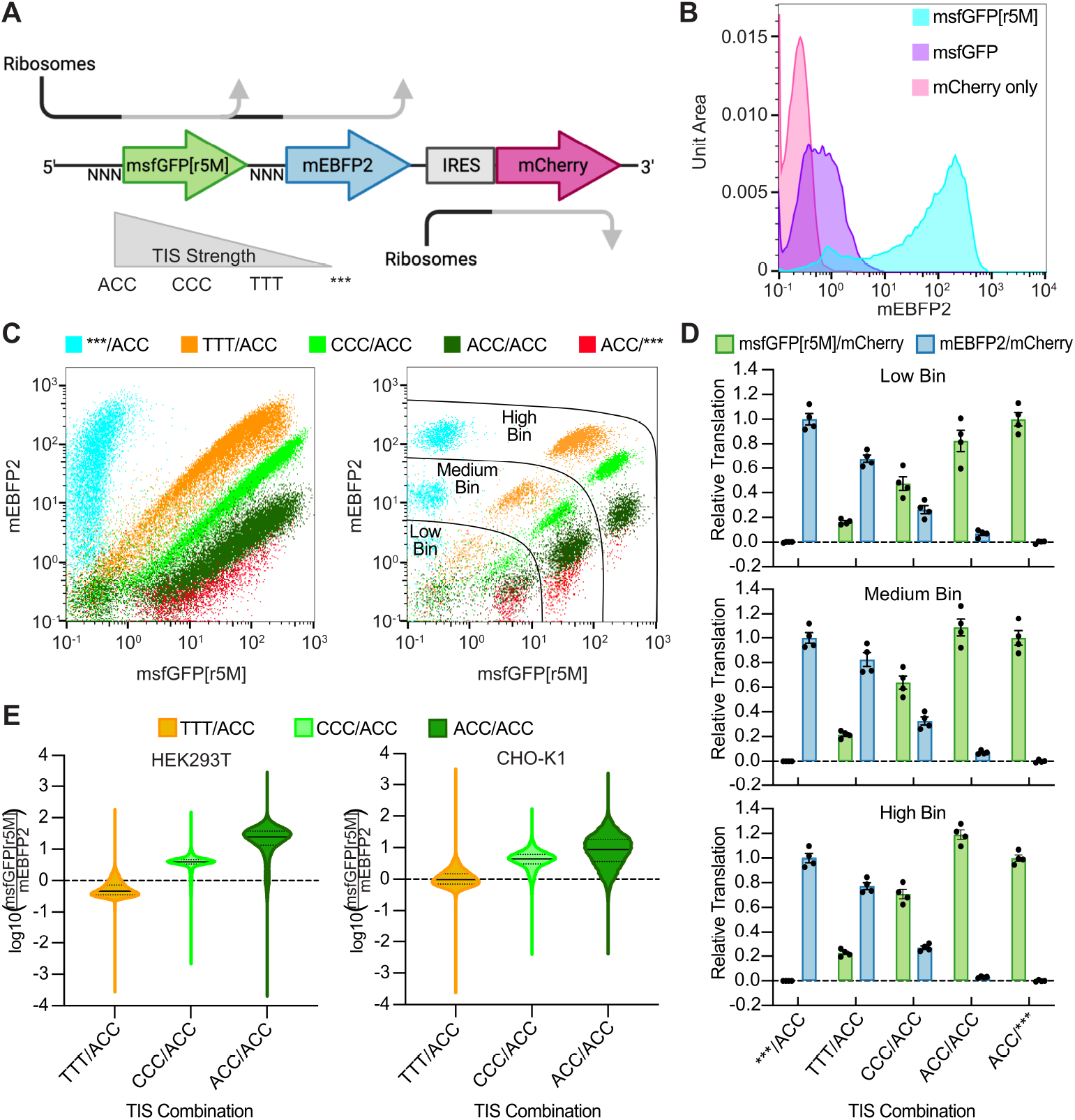
2-ORF SEMPER constructs demonstrate tunable, bicistronic expression. **A**) Architecture of an mRNA transcript produced by transfected 2-ORF SEMPER plasmid DNA. Cap-dependent ribosomes translate ORF 1, msfGFP[r5M], or ORF 2, mEBFP2, with frequencies dependent on the trinucleotide (NNN) upstream of each ORF. The relative strengths of the trinucleotides and TISs they represent are depicted. *** (TTTCCAT) does not contain a start codon, preventing translation of the ORF. IRES-mCherry is included to normalize fluorescence measurements. **B**) mEBFP2 distribution plots for constructs containing different versions of monomeric Superfolder GFP encoded in ORF 1. Removal of internal methionines from the msfGFP leads to much stronger expression of the downstream mEBFP2. Both ORFs used the strong ACC TIS (N=1, representative of four replicates). **C**) Flow cytometry plots of mCherry positive HEK293T cells transfected with 2-ORF SEMPER constructs. The legend contains the ORF 1 TIS and the ORF 2 TIS separated by a slash. (Left) Increasing the strength of the first TIS increases msfGFP[r5M] fluorescence relative to that of mEBFP2 (N=1, representative of four replicates). (Right) Binning cells into three mCherry fluorescence ranges produces unique clusters of cells (N=1, representative of four replicates). **D**) Min-max normalized fluorescence values for each GOI relative to ACC/*** and ***/ACC. Fluorescence measurements were first normalized by mCherry before min-max normalization. Error bars depict standard error of the mean (SEM) (N=4). **E**) Violin plots of log10(msfGFP[r5M]/mEBFP2) values for all mCherry positive cells for four combined replicates of TTT/ACC, CCC/ACC, and ACC/ACC plasmids transfected into HEK293T and CHO-K1 cell lines. The median and quartiles of the distribution are represented by the solid and dotted lines respectively.

After cloning various combinations of TISs in front of our two ORFs, we transfected these “2-ORF SEMPER” constructs into HEK293T cells—a widely used research model for mammalian cell biology. Three single-color control plasmids with msfGFP[r5M], mEBFP2, and mCherry were also transfected. We screened the transfected cells using flow cytometry, utilizing the single-color control plasmids for compensation and correction of fluorescence spillover emissions. As hypothesized, our cell lines produced both msfGFP[r5M] and mEBFP2 from single transcripts (**Figure 1C**). Strikingly, the tested TIS combinations (TIS for ORF 1 / TIS for ORF 2) yielded unique relationships between the fluorescence of the first ORF and that of the second ORF, with the relative expression of the former vs the latter following the strength of the first TIS. To analyze the 2-ORF SEMPER performance as a function of transcript abundance or “copy number”, we binned cells into three categories (low copy, medium copy, and high copy) based on their mCherry fluorescence. This yielded distinct clusters. We then calculated means of msfGFP[r5M]/mCherry, mEBFP2/mCherry, and msfGFP[r5M]/mEBFP2 for the cells in each bin and for each tested TIS combination. We conducted min-max normalization for each ORF separately using two constructs from our screen (***/ACC and ACC/***). We reasoned that ***/ACC would produce the maximum amount of mEBFP2 and the minimum amount of msfGFP[r5M] as all 43S PIC flux should bypass the first ORF and initiate translation at the second. Conversely, we expected ACC/*** to produce the maximum amount of msfGFP[r5M] and the minimum amount of mEBFP2. Conducted for each mCherry bin independently, this analysis enabled us to compare translation levels of a particular FP across various TIS combinations relative to its hypothesized maximum and minimum (**Figure 1D**).

As we increased the TIS strength in front of msfGFP[r5M], we observed increases in msfGFP[r5M] relative translation levels and decreases in mEBFP2 relative translation levels for all mCherry bins. Across mCherry bins, we found that the rank order of relative translation levels for mEBFP2 for our different TIS combinations was conserved. Interestingly, the relative translation of msfGFP[r5M] for the ACC/ACC construct was greater than what was observed for the ACC/*** construct for certain bins. We had originally hypothesized that these constructs would translate nearly the same levels of msfGFP[r5M].

To confirm generalizability across species, we also demonstrated that the 2-ORF SEMPER constructs yielded TIS combination-dependent relative translation levels of our ORFs in CHO-K1 cells, a widely used cell line for the production of biologics (**Figure S1A-B**).^26^ Upon comparing the distributions of log10(msfGFP[r5M]/mEBFP2) values between cell types (**Figure 1E**), we determined that the three TIS combinations tested maintained the same rank order in both HEK293T and CHO-K1 cell lines.

### Scaling plasmid-based SEMPER constructs to three ORFs

Enzymatic pathways and macromolecular assemblies are often composed of more than two proteins or peptide subunits.^2,6,27^ To determine if the SEMPER system could be scaled beyond two ORFs to meet the needs of increasingly complex pathways and assemblies, we screened a variety of “3-ORF SEMPER” constructs. Following a similar strategy to the 2-ORF system, we first cloned and tested a methionine-less monomeric Superfolder BFP (msfBFP[r5M]). We then encoded msfBFP[r5M], msfGFP[r5M], and emiRFP670, a far-red fluorescent protein, into a variety of 3-ORF SEMPER plasmids—maintaining IRES-mCherry for normalization (**Figure 2A**). To conduct min-max normalization, we cloned three plasmids where the starting TISs were removed from two of the ORFs using the *** sequence, while maintaining a strong TIS on the third: i)***/***/ACC, ii) ***/ACC/***, iii) ACC/***/***. The emiRFP670 CDS still encoded methionines within it, yielding some observable far-red fluorescence in constructs ii) and iii). Additionally, we constructed and transiently transfected six other 3-ORF SEMPER plasmids with unique TIS combinations into HEK293T cells and screened them using flow cytometry. Four single-color controls (msfBFP[r5M], msfGFP[r5M], emiRFP670, and mCherry) were also transfected and screened to allow for compensation. Following a similar mCherry binning strategy conducted on the flow cytometry data for 2-ORF SEMPER constructs, we analyzed the mCherry normalized translation levels of our three ORFs relative to their own theoretical maximums and minimums from i), ii), iii) (**Figure 2B**). With the tested TIS combinations, we achieved a wide range of relative translation levels for each ORF, demonstrating tunable co-expression of three genes from a single transcriptional unit.

**Figure 2:**
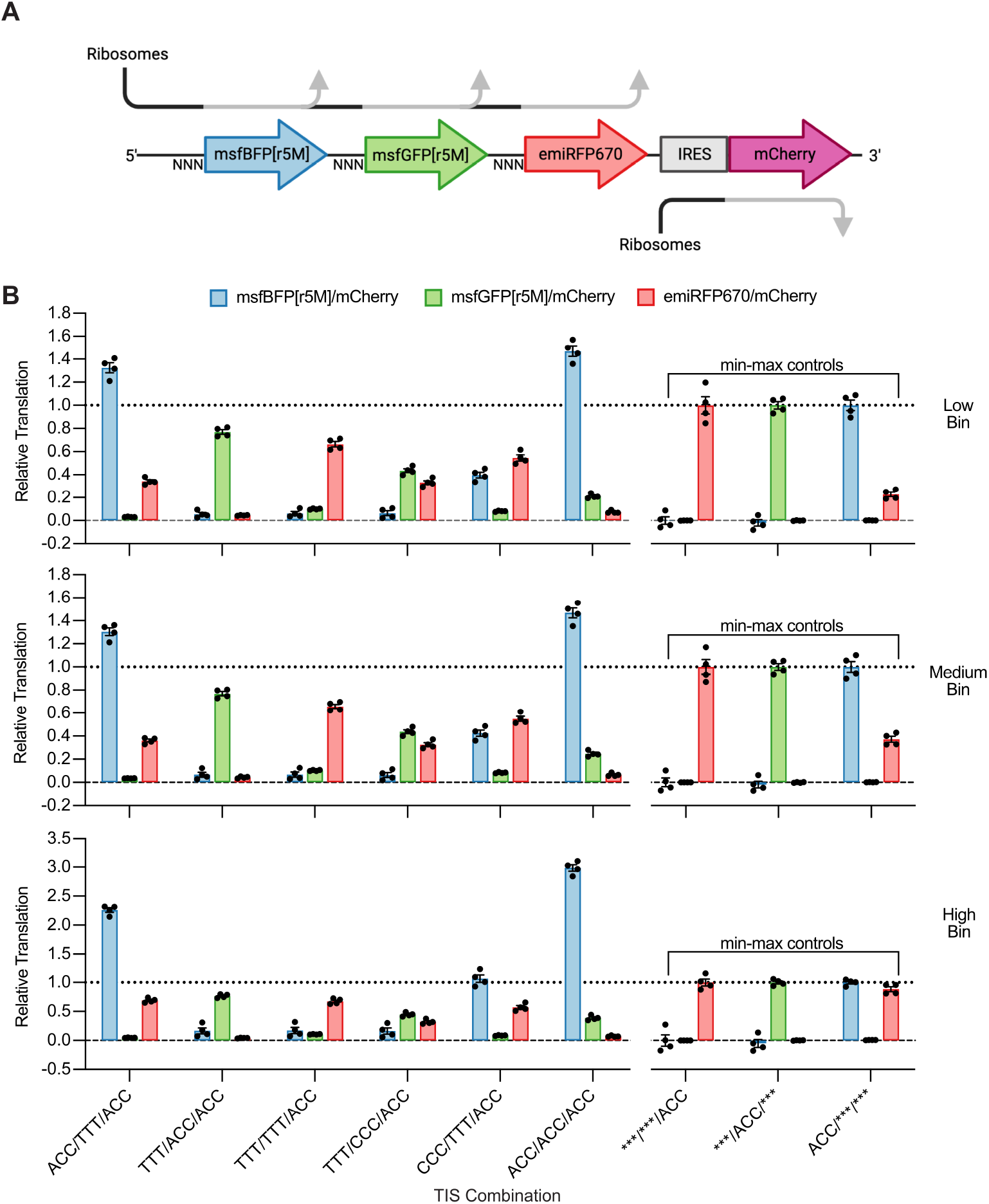
Scaling tunable polycistronic expression beyond two ORFs. **A**) Architecture of an mRNA transcript produced by transfected 3-ORF SEMPER plasmid DNA. **B**) Min-max normalized relative translation levels for each fluorescent protein. Cells were first clustered into three mCherry fluorescence bins (low, medium, high). Min-max controls are depicted on the right of each bar graph. As depicted, *** in front of emiRFP670 yields observable fluorescence for some constructs as emiRFP670 contains methionines within its sequence, allowing for translation of partial peptides with fluorescence activity. Error bars depict SEM (N=4).

Notably, two combinations, ACC/TTT/ACC and ACC/ACC/ACC, yielded relative translation levels for msfBFP[r5M] greater than 1.0 in HEK293T cells across all mCherry bins. We also observed values for msfBFP[r5M] much greater than 1.0 in CHO-K1 cells for a majority of TIS combinations in both medium and high bins (**Figure S2A**). Wu et al. previously reported that the presence of downstream ORFs can increase the translation of upstream ones.^28^ We hypothesize that constructs containing multiple ACC TISs have a higher prevalence of translating ribosomes, allowing these ribosomes and subunits—with their documented helicase activity—to maintain the mRNA secondary structure in an open conformation that is more amenable for ribosomal scanning, translation initiation, and subsequent protein expression.^29^ However, more work is required to uncover the exact mechanisms that elicit this observed phenotype. For multiple TIS combinations, we found that the second and third ORFs achieved relative translation levels greater than 50% while still producing substantial levels of upstream ORF products. These results suggest there may be sufficient ribosomal flux to effectively scale the SEMPER paradigm to 4+ ORFs.

### Producing gas vesicle ultrasound reporters using 2-ORF SEMPER

Next, we set out to demonstrate that 2-ORF SEMPER constructs could be applied to encoding a multimeric protein complex by expressing gas vesicles (GV) using mammalian acoustic reporter genes (mARGs).^30^ Originally evolved in prokaryotes, GVs were recently introduced as genetically-encodable reporters for ultrasound imaging, enabling the noninvasive imaging of dynamic cellular processes in living organisms.^2,31^ A single GV is made up of many GvpA structural units that are assembled together in a helical pattern through the cooperative activity of six heterologous assembly factors and minor constituents, referred to collectively as GvpNJKFGW.^32,33^ Using a two-vector system, our group has previously expressed GVs in mammalian cells by co-transfecting one plasmid encoding the structural unit upstream of IRES-mCherry (pgvpA-IRES-mCherry) along with another plasmid encoding the assembly factors and a terminal Emerald GFP (EmGFP), all linked together by P2A elements (pgvpNJKFGW-EmGFP). Additionally, we have found that co-transfecting the pgvpA-IRES-mCherry in excess of pgvpNJKFGW-EmGFP improves acoustic contrast.^30^ Likewise, *Anabaena flos-aquae*—the organism from which mARGs used in this study are derived—contains more copies of gvpA relative to the other gvps in its GV gene cluster.^34^

Using the 2-ORF SEMPER strategy, we cloned single-vector systems for producing GVs in mammalian cells. We refer to these constructs as SEMPER mARGs. As gvpA does not contain any internal methionines, we encoded it directly into the first ORF. We tested our panel of TIS sequences in front of gvpA while maintaining the strong ACC TIS in front of the gvpNJKFGW-EmGFP ORF (**Figure 3A**). We compared these SEMPER constructs to our published two-vector expression system, mixing the gvpA-IRES-mCherry plasmid in 4-fold molar excess of the gvpNJKFGW-EmGFP plasmid. We transfected these plasmids into HEK293T cells, maintaining the same total mass of plasmid for each transient transfection. As the strength of the TIS in front of gvpA increased, we found that the translation level of gvpNJKFGW-EmGFP, as measured by Emerald/mCherry, decreased (**Figure 3B**), as expected from our 2-ORF SEMPER FP experiments. As measured by BURST ultrasound imaging,^35^ the ACC/ACC combination—predicted to yield the highest ratio of GvpA to GvpNJKFGW— produced the strongest acoustic contrast compared to the other SEMPER mARG plasmids (**Figure 3C-D**).

**Figure 3:**
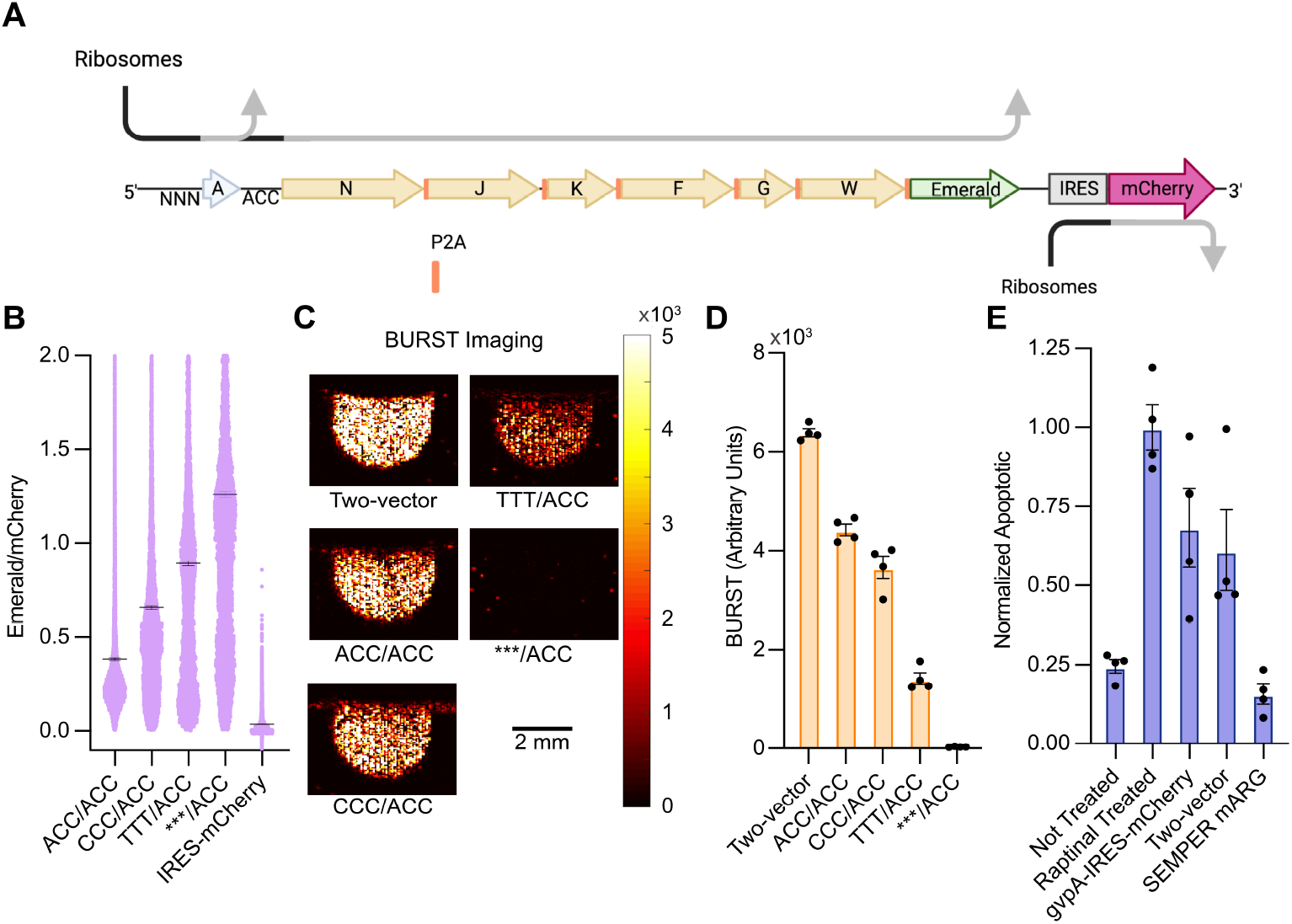
Utilizing 2-ORF SEMPER constructs to express gas vesicles in mammalian cells. **A**) Architecture of an mRNA transcript transcribed from transfected SEMPER mARG plasmid DNA. The first ORF, gvpA, encodes the main structural protein, while the second ORF, gvpNJKFGW-EmGFP, encodes all necessary accessory proteins strung together with P2A self-cleaving peptides. **B**) Flow cytometry distributions of Emerald GFP normalized by mCherry values for each TIS combination tested in addition to an IRES-mCherry control. The thick line depicts the mean of the distribution. Error bars depict standard error of the mean (SEM) (N=1, representative of four replicates). **C**) BURST images of acoustic contrast due to gas vesicle expression within HEK293T cells. Depicted are HEK293T cells loaded into agarose phantoms three days after transfection of SEMPER mARG plasmids or the leading two-plasmid system. The color bar represents the magnitude of BURST signal measured in linear arbitrary units. The floor and ceiling of the images are set to 0 and 5000 respectively (N=1, representative of four replicates). **D**) BURST signal quantification of gas vesicle acoustic contrast for HEK293T samples transfected with SEMPER mARG plasmids or the leading two plasmid system. Error bars depict SEM (N=4). **E**) Annexin V staining assays to quantify the number of apoptotic cells following expression of mARG vectors. HEK293T cells transfected with pgvpA-IRES-mCherry with the start codon removed in front of gvpA were used to establish a baseline (Not Treated). A subset of these cells was treated with Raptinal to induce apoptosis (Raptinal Treated). Other cell populations were transfected solely with a fully functional pgvpA-IRES-mCherry (gvpA-IRES-mCherry), the two-vector system (Two-vector), or the ACC/ACC SEMPER mARG plasmid (SEMPER mARG). Error bars depict SEM (N=4).

### SEMPER mARG expression system reduces cell toxicity

A significant issue with multimeric protein assemblies is the potential for cellular burden or toxicity due to imperfect stoichiometry or the absence of an essential assembly component or chaperone. In our experiments with mARGs, we observed that samples transfected with the two-vector system contained a larger fraction of cells that were positive for pgvpA-IRES-mCherry but negative for pgvpNJKFGW-EmGFP (39.03% ±1.42%) compared to cells transfected with SEMPER mARG plasmids (13.40%±0.85%) (**Figure S3A**). This is not surprising due to the inherent stochasticity of transient co-transfection. As GvpA subunits have been speculated to nonspecifically aggregate when expressed without GV assembly factors,^36,37^ we hypothesized that cells receiving a sub-optimal ratio of pgvpA-IRES-mCherry : pgvpNJKFGW-EmGFP or only pgvpA-IRES-mCherry may have higher incidences of apoptosis due to the formation of cytotoxic GvpA aggregates in the cytoplasm.

To test this hypothesis, we performed Annexin V-based apoptosis assays on cells transfected with either pgvpA-IRES-mCherry alone, the two-vector mARG expression system at optimal transfection ratio, or the ACC/ACC SEMPER mARG (**Figure 3E**). In each condition, molar amounts of gvpA gene were equalized in each transfection mixture. In addition, pgvpA-IRES-mCherry plasmid without a start codon in front of gvpA was transfected into HEK293T cells to establish a negative control. A subset of these samples was then treated with Raptinal to induce apoptosis, establishing a positive control. Notably, expressing GvpA without its assembly factors led to high levels of apoptosis.

While co-transfecting pgvpNJKFGW-EmGFP reduced some of the observed toxicity, the ACC/ACC SEMPER mARG transfected cells were significantly healthier, with apoptosis levels indistinguishable from untreated negative controls. Taken together, these results suggest that the SEMPER mARG plasmid reduces cell toxicity by ensuring that assembly factors are consistently co-expressed with structural proteins in an appropriate ratio.

### Demonstrating SEMPER 2-ORF efficacy from transfected synthetic mRNA

With accelerating advancements in mRNA vaccine technology and growing development efforts in mRNA-based therapeutics,^10,38^ we sought to demonstrate the efficacy of the SEMPER 2-ORF system to tune relative protein expression stoichiometries using mRNA-based expression systems (**Figure 4A**). Our *in vitro* transcribed (IVT) mRNA constructs do not include IRES-mCherry, as we observed reduced expression of our GOIs in IVT mRNA constructs containing this element (**Figure S4**).

**Figure 4:**
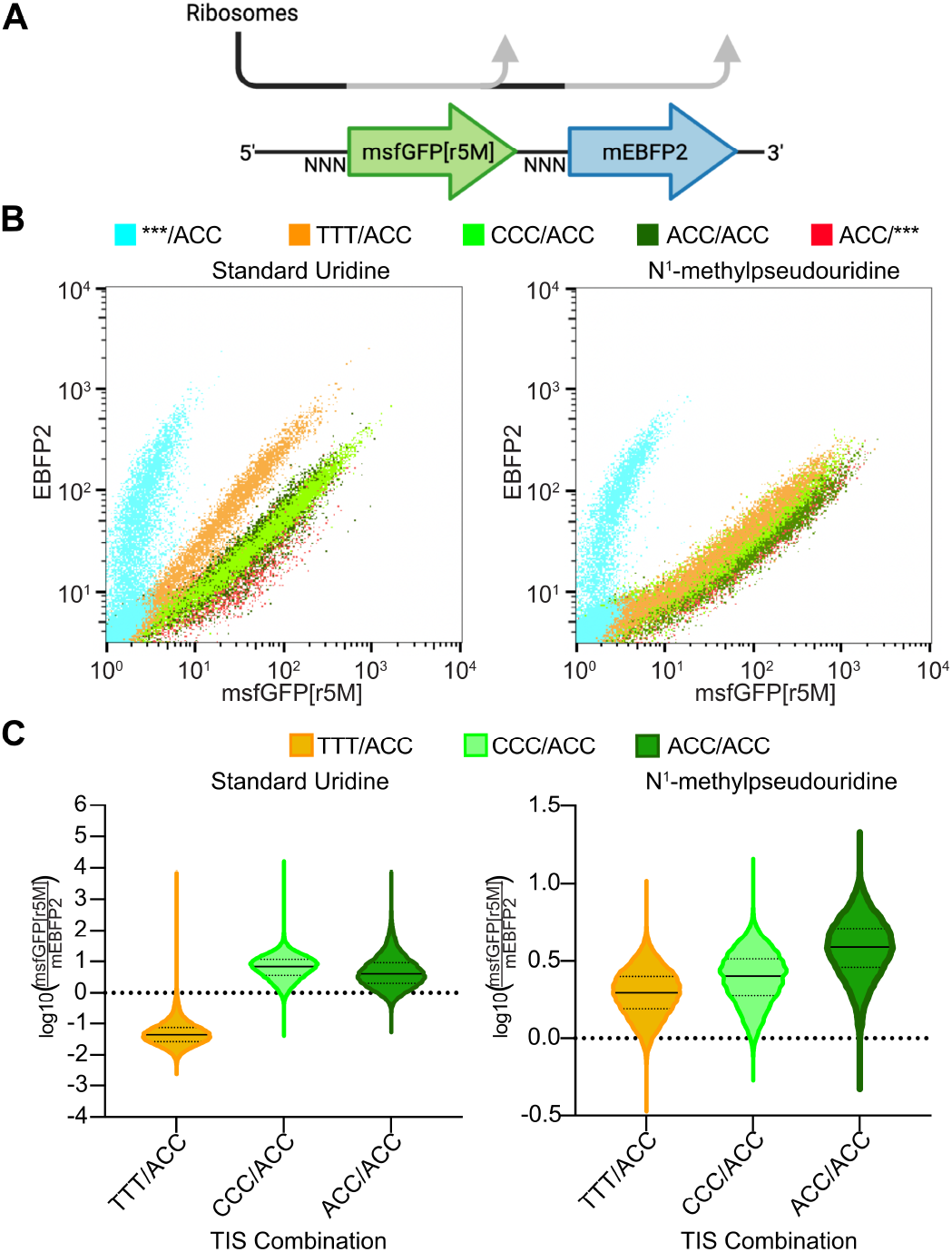
2-ORF SEMPER using *in vitro* transcribed mRNA. **A**) Schematic of 2-ORF SEMPER mRNA produced from *in vitro* transcription. The first and second ORFs encode msfGFP[r5M] and mEBFP2 respectively. **B**) Flow cytometry data from transient transfection of IVT mRNA encoding various TIS combinations into HEK293T cells (N=1, representative of four replicates). IVT mRNA constructs were made with (Left) standard uridine triphosphate or (Right) N^1^-methylpseudouridine-5’-triphosphate. **C**) Violin plots of log10(msfGFP[r5M]/mEBFP2) values for msfGFP[r5M] and mEBFP2 double-positive cells transfected with TTT/ACC, CCC/ACC, or ACC/ACC IVT mRNA. Each distribution represents four combined replicates. The median and quartiles of the distribution are represented by the solid and dotted lines respectively.

Through IVT and subsequent purification steps, we produced concentrated mRNA samples with the same TIS combinations and ORFs as tested for the plasmid-based 2-ORF SEMPER constructs. As nucleotide chemistry has been shown to impact translation efficiency,^39^ we hypothesized that changing the nucleotide chemistry of our IVT mRNA may lead to changes in expression of our two ORFs. Therefore, we created a set of mRNA samples using standard uridine triphosphate (UTP) nucleotides in the IVT reaction, while another set was made using N^1^-methylpseudouridine-5’-triphosphate (m1Ѱ), commonly used to manufacture mRNA for clinical use.^40^ We then transiently transfected these mRNA samples into HEK293T cells and screened them using flow cytometry. Additionally, we transfected and collected flow cytometry data for two plasmid-based single-color controls for compensation.

Flow cytometry showed that some TIS combinations yielded unique relationships between msfGFP[r5M] and mEBFP2 fluorescence (**Figure 4B-C**). When using standard UTPs, the relationship for TTT/ACC was significantly different than those of CCC/ACC and ACC/ACC, demonstrating that multiple product stoichiometries are accessible from IVT mRNA SEMPER constructs. However, the relationships for CCC/ACC and ACC/ACC were not as distinct as those achieved at the plasmid level, suggesting that further IVT-specific tuning is required. In comparison, the mEBFP2 vs msfGFP[r5M] relationships for TTT/ACC, CCC/ACC, and ACC/ACC mRNA constructs produced with m1Ѱ were all very similar, confirming that nucleotide chemistry impacts translation initiation.

## DISCUSSION

Our results demonstrate that SEMPER enables tunable expression of multiple recombinant ORFs from single transcripts in mammalian cells by using leaky ribosomal scanning and variable strength TISs. The SEMPER paradigm is applicable across cells from multiple species and can yield a range of user-tunable expression stoichiometries for at least three GOIs. By demonstrating GV production with SEMPER mARGs, we highlight that this novel system for polycistronic expression can be used to encode complex multi-gene constructs and titrate the relative stoichiometries of translated products to optimize expression and minimize toxicity in mammalian cells. Finally, we show the SEMPER framework can be applied directly to mRNA-based genetic circuits by demonstrating polycistronic expression with 2-ORF SEMPER mRNA constructs using IVT. Furthermore, we show that nucleotide chemistry can alter the translation levels of encoded ORFs in IVT mRNA, opening new avenues to further tune protein stoichiometries with synthetic mRNA.

Although the genetic constructs described in this work should be immediately useful in a variety of contexts, future work is needed to demonstrate the SEMPER framework’s utility upon integration into the genome through stable transfection and viral transduction methods. Another important future step is to make SEMPER compatible with methionine-containing proteins. Internal methionine codons create TISs that may undesirably consume ribosomal flux from downstream open reading frames. If mutating these methionines is not possible, an alternative approach may be to alter the nucleotide context surrounding the in-frame AUG using rationally chosen degenerate codons to effectively mask the TIS.

We believe SEMPER represents a significant advancement in the field of recombinant multi-protein expression and synthetic biology, enabling user-friendly tuning of multiple proteins within mammalian systems. Researchers can readily adopt this framework to rapidly screen libraries relevant to expression and tuning of enzymatic pathways and circuits, assembly of multimeric protein structures, and other endeavors.^41^ As we have demonstrated in this study, this technology can enable simultaneous expression of a protein of interest and its folding chaperones from a single transcriptional unit at an optimal ratio, offering a novel strategy to tackle protein misfolding precisely. SEMPER’s compact framework and tunability make it a critical step towards increasingly complex engineering efforts and finer control of mammalian systems. Moreover, as shown in some of the constructs used in this study, SEMPER can be used in combination with IRES and 2A elements for versatile encoding of more complex genetic constructs.

SEMPER also holds potential in the field of RNA vaccines and therapeutics. It could enhance the development of polyvalent RNA vaccines as well as the expression of mosaic virus-like particles from delivered mRNA, thereby broadening the immune response against highly variable viruses.^42,43^ Further, it could improve RNA-delivered monoclonal antibody therapeutics by optimizing the ratio of heavy and light chain production in each cell, and by allowing for simultaneous production of multiple or bi-specific antibodies from a single mRNA.^44–46^ This technology also paves the way for the production of cytokine cocktails through the co-expression of multiple cytokines from a single mRNA, which could offer synergistic effects to modulate immune responses. Taken together, SEMPER provides an option that can always be considered in developing polycistronic constructs for synthetic biology and medicine.

## MATERIALS AND METHODS

### Vector construction

Plasmids were constructed using standard cloning techniques, including Gibson assembly and conventional restriction and ligation. All final sequences were verified using whole-plasmid sequencing through Primordium Labs. Plasmids containing two or three fluorescent protein ORFs were constructed in the following way: Individual FP sequences were ordered from IDT or TWIST Biosciences as synthetic gBlocks. Modified TIS sequences and the *** sequence were introduced using overhang PCR primers. Finally, all components were subcloned into pCMV-Sport-gvpA-IRES-mCherry using NEB HiFi DNA Assembly replacing the gvpA ORF. Some modifications were made to the 5’ UTR sequences. Plasmid sequences for the 2-ORF TTT/ACC construct and the 3-ORF ACC/***/*** construct as well as sequences for individual genetic parts in these plasmids are provided in **Table S1** to aid in reproduction of this work.

SEMPER mARG plasmids containing gvpA and gvpNJKFGW-EmGFP were constructed as follows: The ORF containing gvpNJKFGW-EmGFP was PCR amplified and subcloned into pCMV-Sport-gvpA-IRES-mCherry using NEB HiFi DNA Assembly between gvpA and IRES. Different gvpA TIS sequences were introduced using single-stranded oligo bridge primers with NEB HiFi Assembly.

To produce plasmids amenable for *in vitro* transcription of mRNA, sequences containing the 5’UTR through the stop codon of the mEBFP2 ORF from the 2-ORF SEMPER plasmids were cloned into an IVT plasmid backbone containing a T7 promoter, a synthetic 3’UTR sequence, and a 100 bp polyA track (**Table S1**) using PCR amplification and Gibson assembly methods.

### Production of mRNA by *in vitro* transcription

Linear templates were PCR amplified from the IVT plasmids (described in Vector construction) using primer p101 and “Ultramer” p54 synthesized by Integrated DNA Technologies. Primer p101 introduced an AG dinucleotide following the T7 promoter sequence on linear templates to ensure efficient 5’ capping of IVT mRNA. Linear templates were purified using standard gel electrophoresis and DNA cleanup methods. IVT mRNA was produced using the HiScribe T7 High Yield RNA Synthesis Kit (NEB # E2040) along with 500 ng of linear template and 4 mM CleanCap AG Reagent (Trilink, N-7113). 5 mM N^1^-Methylpseudouridine-5’-triphosphate (Trilink, N-1081) was substituted for the standard uridine triphosphate included in the HiScribe Kit for certain reactions. Following the IVT incubation, reactions were additionally incubated with 2 units of DNAse I to remove remaining DNA template. Purification of IVT mRNA was conducted using the Monarch RNA Cleanup Kit (NEB #T2040L). To confirm IVT mRNA were of the correct lengths, aliquots of purified IVT mRNA were denatured at 70°C for 15 minutes and then subjected to gel electrophoresis on a 1% agarose gel in Tris acetate EDTA (TAE) stained with SYBR Safe. ssRNA ladder (NEB #N0362S) was used to confirm band sizes. mRNA concentrations for each IVT mRNA were quantified using a Qubit Fluorometer (Invitrogen). Completed IVT mRNA samples were stored at -80°C.

### Cell Culture and Transfection

HEK293T cells (American Type Culture Collection (ATCC), CLR-2316) and CHO-K1 cells (American Type Culture Collection (ATCC), CCL-61) were cultured in 24-well plates at 37 °C and 5% CO2 in a humidified incubator in 0.5 mL of DMEM (Corning, 10-013-CV) with 10% FBS (Gibco) and 1× penicillin–streptomycin until about 80% confluency before transfection with plasmid DNA or IVT mRNA.

For plasmid DNA transfection, transient transfection mixtures were created by mixing 500 ng of plasmid with polyethyleneimine (PEI-MAX, linear 40 kD #24765-2, Polysciences) at 4.12 μg of polyethyleneimine per microgram of DNA in 150 mM NaCl. The mixture was incubated for 12 minutes at room temperature and added dropwise to HEK293T or CHO-K1 adherent cells. Media was changed after 12–16 hours and daily thereafter. Cells were analyzed with flow-cytometry 48 hours post-transfection.

For IVT mRNA transfection, transient transfection mixtures were created by mixing 500 ng of mRNA, 1.5 μL of Lipofectamine MessengerMAX Reagent (Invitrogen #LMRNA008), and 50 μL of Opti-Mem media (Gibco #31985070) according to the Lipofectamine MessengerMAX Reagent standard operating procedure. The mixture was incubated for 5 minutes at room temperature and added dropwise to HEK293T cells. Media was changed after 12–16 hours and daily thereafter. Cells were analyzed with flow-cytometry 48 hours post-transfection.

For transfection of plasmids encoding mARGs, transient transfection mixtures were created as above except that 600 ng of total DNA was mixed together as follows: 56 fmol of gvpA-IRES-mCherry plasmid with or without 14 fmol gvpNJKFGW-EmGFP plasmid or 56 fmol of a 2-ORF SEMPER mARG plasmid. Plasmid mixtures were normalized with addition of pUC19 up to 600 ng before complexing with PEI-MAX. The mixture was incubated for 12 minutes at room temperature and added dropwise to HEK293T adherent cells. Media was changed after 12–16 hours and daily thereafter. Cells were harvested 3-days post-transfection.

### Flow Cytometry and Apoptosis Assays

To harvest the cells, cells were dissociated using trypsin/EDTA and centrifuged at 300 g for 6 minutes at room temperature. For experiments involving mARGs, 2-ORF SEMPER plasmid transfections, and 2-ORF SEMPER mRNA transfections, cells were analyzed with MACSQuant VYB (Miltenyi Biotec). For experiments involving 3-ORF SEMPER, cells were analyzed using Cytoflex S (Beckman Coulter). Single-color controls were used to allow for compensation. In plasmid transfection experiments, cells were gated for size and doublet-discriminated before being gated and further binned by mCherry fluorescence. For mRNA transfection experiments cells were gated as above except they were gated for msfGFP[r5M] and mEBFP2 double-positivity instead of by mCherry fluorescence.

For the Annexin V apoptosis assay, we transfected cells as described above but did not perform media changes. For the positive control, cells transfected with pgvpA-IRES-mCherry (w/ start codon removed in front of gvpA) were treated with 10 μM Raptinal 18 hours before harvest. Cells were harvested two days after transfection. To do this, we collected supernatant from cells to recover the dead cells. We then trypsinized the adhered cells and added this trypsinized cell fraction to the original supernatant for maximum cell recovery. Then, we spun the cells down at 300 g for 6 minutes, carefully removed the supernatant, and resuspended the cells in 90 μL of cold annexin binding buffer with no EDTA and supplemented Ca^2+^, followed by 10 μL of Annexin V Pacific Blue stain (A35122). Following a 15 minute incubation, we added 150 μL of the same annexin binding buffer and then performed flow cytometry to obtain fluorescence values for the pacific blue stain. We gated in FlowJo by selecting the largest population in the FCS/SSC plot, keeping the lower left corner in the cell gate to include smaller apoptotic cells. We selected a 10^4^ threshold for the Pacific Blue Annexin V stain, corresponding to the right tail of the unstained control so that the Annexin V+ population of the unstained control was around 0%. Values were normalized by setting the mean value of Raptinal-treated samples to 100%.

### In vitro ultrasound imaging of transient expression of GVs in HEK293T cells suspended in agarose phantoms

Cells were harvested as described above. Cells were resuspended with 1% low-melt agarose (GoldBio) in PBS at 40°C at concentrations of ∼15 million cells per milliliter and then loaded into wells of pre-formed phantoms consisting of 1% molecular biology-grade agarose (Bio-Rad) in PBS. Phantoms were imaged using L22-14vX transducer (Verasonics) at 15.625 MHz while submerged in PBS on top of an acoustic absorber pad. For BURST imaging, wells were centered around the 8 mm natural focus of the transducer and a BURST pulse sequence was applied in pAM acquisition mode with the focus set to 8 mm, and the voltage was set to 2V for the first 10 frames and 15V for the remaining frames.

BURST images were produced by pixel-wise subtraction of the 11th (collapse) frame from the 54th (post-collapse) frame. Resulting differences were divided by 100. Images were quantified as follows: The sample ROIs were drawn inside the well of the agarose phantom. Average pixel value inside the ROI was calculated for each replicate.

## Supporting information

All Supplementary Information

## ACKNOWLEDGEMENTS

The authors thank Michael Elowitz for helpful discussions and James Linton for equipment training. This work was supported by the National Institutes of Health (R01EB018975 to MGS), the Millard and Muriel Jacobs Genetics and Genomics Laboratory at the California Institute of Technology, and the Flow Cytometry and Cell Sorting Facility at California Institute of Technology. Related research in the Shapiro laboratory is supported by the Chan Zuckerberg Initiative. MGS is an investigator of the Howard Hughes Medical Institute. Some graphics were created with BioRender.com.

