## Supplementary material for "Stoichiometric expression of messenger polycistrons by eukaryotic ribosomes (SEMPER) for compact, ratio-tunable multi-gene expression from single mRNAs": All Supplementary Information

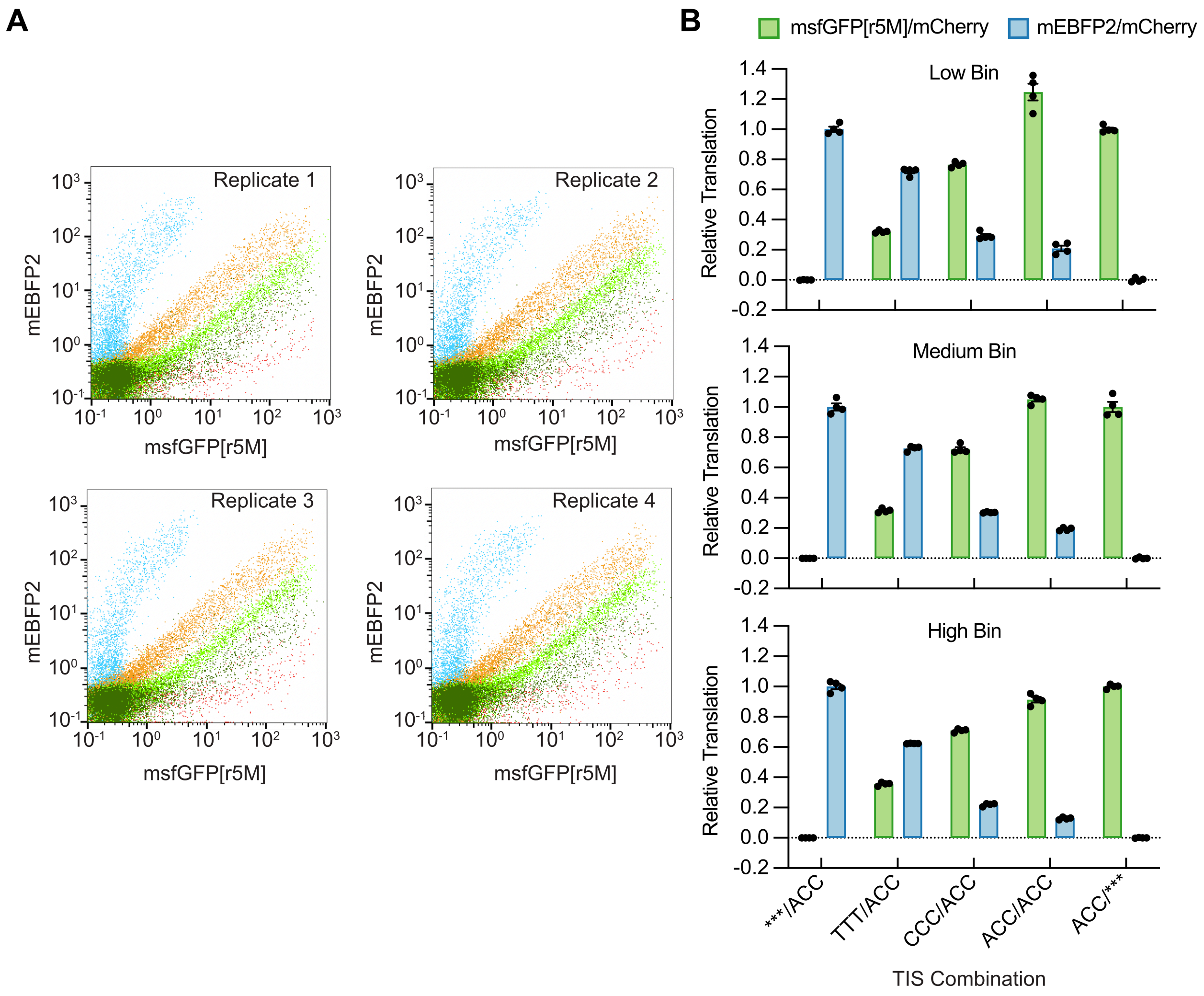


**Figure S1: CHO-K1 plasmid-based 2-ORF SEMPER results A**) Flow cytometry data for CHO-K1 cells transfected with 2-ORF SEMPER constructs. mCherry positive cells are depicted. Transfection conditions were optimized for HEK293T cells and therefore lower transfection efficiency was observed for CHO-K1 cells. Replicates for both cell lines demonstrated reproducibility as shown here for CHO-K1. **B**) Min-max normalized fluorescence values for each GOI relative to ACC/*** and ***/ACC. Fluorescence measurements were first normalized by mCherry before min-max normalization. Error bars depict SEM (N=4).

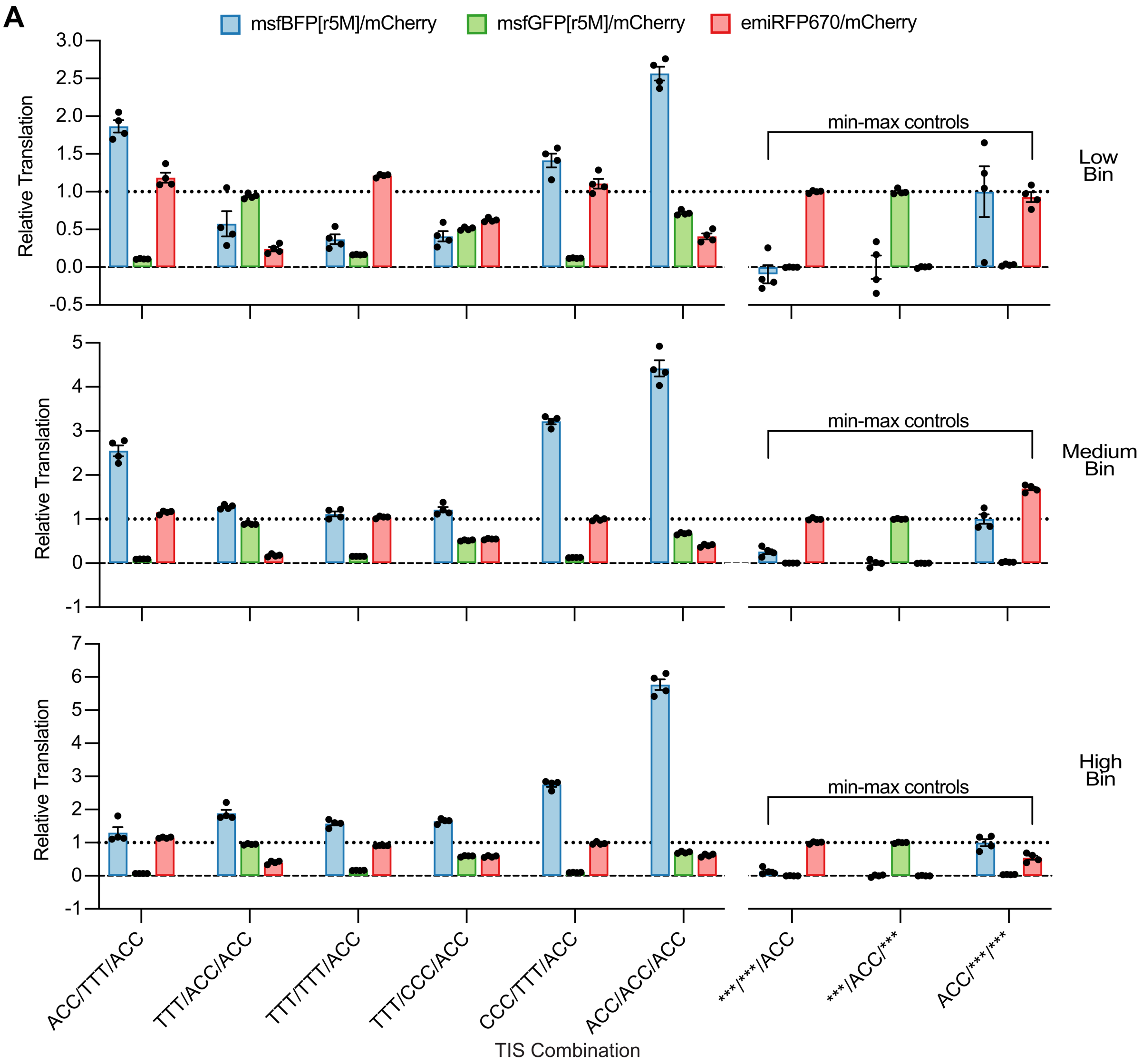


**Figure S2: 3-ORF SEMPER constructs in CHO-K1 cells. A**) Min-max normalized relative translation levels for each fluorescent protein. Cells were first clustered into three mCherry fluorescence bins (low, medium, high). Constructs used for min-max calculations are depicted on the right of each bar graph. As depicted, *** in front of emiRFP670 yields observable fluorescence for some constructs as emiRFP670 contains methionines within its sequence, allowing for translation of partial peptides with fluorescence activity. Error bars depict SEM (N=4).

**A**

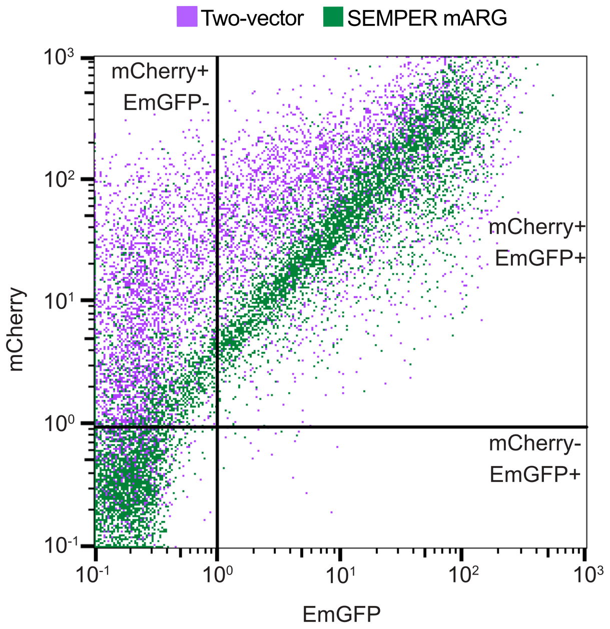


**Figure S3: SEMPER mARGs are not inhibited by the stochasticity of co-transfection of two plasmids**. **A**) Comparison of mCherry and EmGFP fluorescence in HEK293T cells transfected with the two-vector mARG expression system or the SEMPER mARG expression system. The gvpA-IRES-mCherry plasmid was transfected in a 4-fold molar excess relative to pgvpNJKFGW-EmGFP, leading to a significant portion of cells with solely mCherry positivity compared to those transfected with the ACC/ACC SEMPER mARG construct—which contains both fluorescent proteins on a single vector.

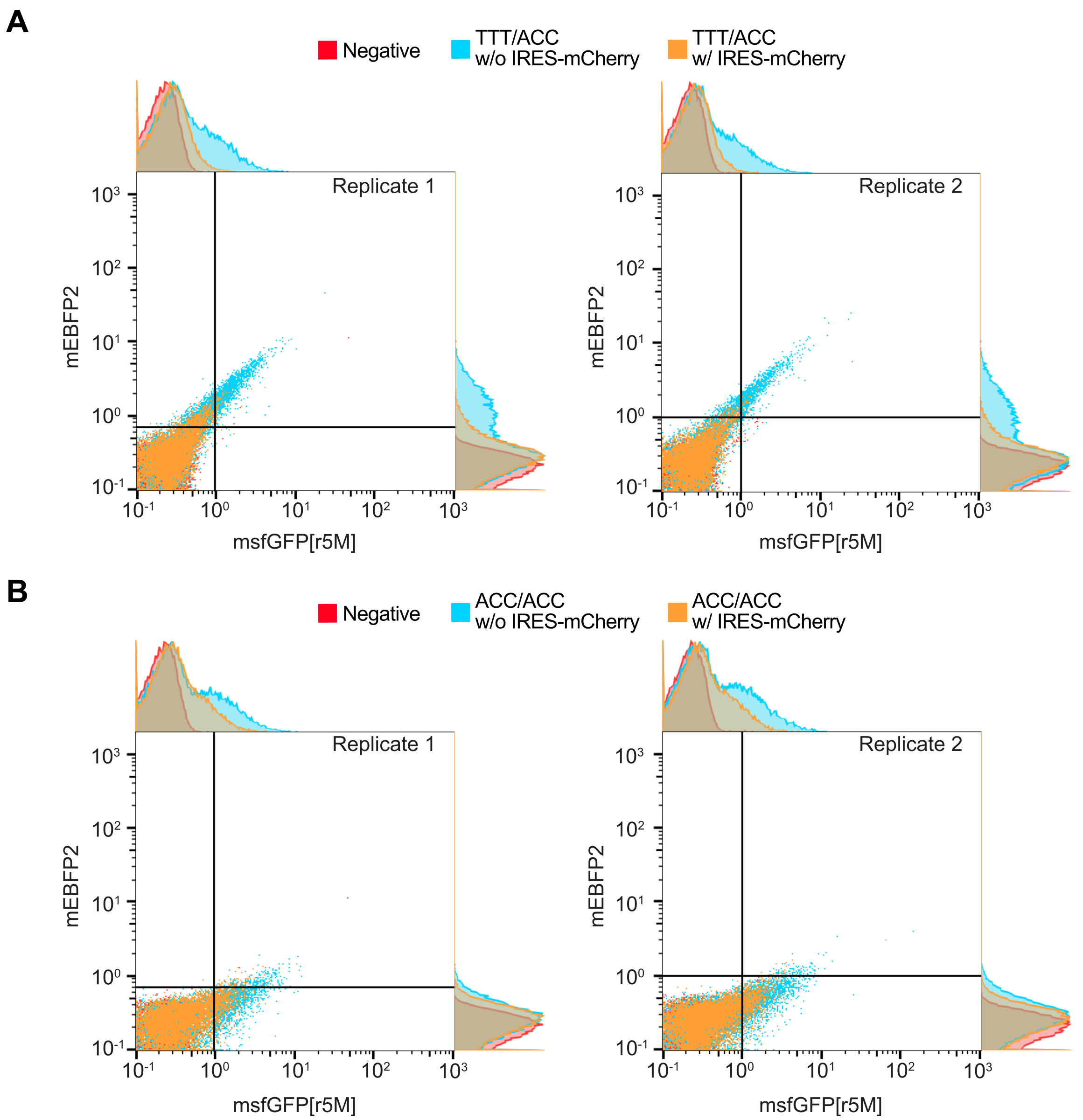


**Figure S4: 2-ORF IVT mRNA tests with and without IRES-mCherry as measured by flow cytometry. A**) TTT/ACC and **B**) ACC/ACC IVT mRNA constructs were synthesized with and without IRES-mCherry. The IVT mRNA tested was produced with standard UTPs. Negative control cells were transfected solely with Lipofectamine MessengerMAX reagent without mRNA. Cells were first gated by size and then doublet-discriminated. Gates drawn above are based on background fluorescence intensities measured for the negative control cells. For both TIS combinations, the IRES-mCherry mRNA constructs yielded lower mEBFP2 and msfGFP[r5m] expression following transfection into HEK293T cells (N=2). IRES-mCherry was therefore not included in IVT mRNA constructs tested and depicted in the main text.

**Table S1: Genetic parts, primers, and plasmid sequences used to construct plasmids and IVT mRNA.**

| Sequence Type | Sequence Identifier | Description | Sequence (5' → 3') |
|---|---|---|---|
| Primer | p101 | Forward primer to generate linear IVT templates. Additionally, this primer ensures there is an AG dinucleotide at the beginning of the transcript. | CGGCCGCTAATACGACTCACTATAAGGG TCAGATCGC |
| Ultramer | p54 | Reverse primer to generate linear IVT templates. This primer ensures the linear template contains a 120 bp polyA tail. | TTTTTTTTTTTTTTTTTTTTTTTTTTTTTTTT TTTTTTTTTTTTTTTTTTTTTTTTTTTTTTTT TTTTTTTTTTTTTTTTTTTTTTTTTTTTTTTT TTTTTTTTTTTTTTTTTTTTTTTTGCTAGCT CCAGGGTGTGG |
| 5' Untranslated Region | 5'UTR (for 2-ORF and 3-ORF plasmids for FP expression) | The 5'UTR used for 2-ORF and 3-ORF plasmids and mRNA SEMPER constructs employing only fluorescent proteins. The trinucleotide NNN that changes the TIS of the first ORF is not included in this 5'UTR | gtcagatcgcctggagacgccatccacgctgttttgacctcc atagaagacaccgggaccgatccagcctccggactctag cctaggcttttgcaaaaagctatttaggtgacactatagaag gtacgcctgcaggtaccgagctcggatccagtagtttaaac cccttc |
| 5' Untranslated Region | 5'UTR for SEMPER mARG plasmids | The following 5'UTR was used for SEMPER mARG plasmids | gtcagatcgcctggagacgccatccacgctgttttgacctcc atagaagaccgagctcggatccagtacccttc |
| 3' Untranslated Region | 3'UTR (plasmid) | The 3'UTR for plasmid-based expression notably includes a Woodchuck Hepatitis Virus Post-transcriptional Regulatory Element to extend transcript half-life, and an hGH polyA signal. | aggatccctcgagggcccaagcttacgcgtgcatgcgac gtcatagctctctccctatagtgagtcgtattataagctagctt gggatctttgtgaaggaaccttacttctgtggtgtgacataatt ggacaaactacctacagagatttaaagctctaaggtaaata taaaatttttaagtgtataatgtgttaaactagctgcatatgctt gctgcttgagagttttgcttactgagtatgatttatgaaaatatt atacacaggagctagtgattctaattgtttgtgtattttagattc acagtcccaaggctcatttcaggcccctcagtcctcacagtc tgttcatgatcataatcagccataccacatttgtagaggtttta cttgctttaaaaaacctcccacacctccccctgaacctgaaa cataaaatgaatgcaattgttgttgttcgtggctgaatcaacct ctggattacaaaatttgtgaaagattgactggtattcttaacta tgttgctccttttacgctatgtggatacgctgctttaatgcctttgt atcatgctattgcttcccgtatggctttcattttctcctccttgtata aatcctggttgctgtctctttatgaggagttgtggcccgttgtca ggcaacgtggcgtggtgtgcactgtgtttgctgacgcaacc cccactggttggggcattgccaccacctgtcagctcctttccg ggactttcgctttccccctccctattgccacggcggaactcat cgccgcctgccttgcccgctgctggacaggggctcggctgt tgggcactgacaattccgtggtgttgtcggggaaatcatcgt cctttccttggctgctcgcctgtgttgccacctggattctgcgc gggacgtccttctgctacgtcccttcggccctcaatccagcg gaccttccttcccgcggcctgctgccggctctgcggcctcttc cgcgtcttcgccttcgccctcagacgagtcggatctccctttg ggccgcctccccgccagagacaattaacttcgcgggtggc atccctgtgacccctccccagtgcctctcctggccctggaag ttgccactccagtgcccaccagccttgtcctaataaaattaa gttgcatcattttgtctgactaggtgtccttctataatattatggg gtggaggggggtggtatggagcaaggggcaagttgggaa |

| | | | |
|---|---|---|---|
| | | | gacaacctgtagggcctgcggggtctattgggaaccaagc<br>tggagtgcagtggcacaatcttggctcactgcaatctccgcc<br>tcctgggttcaagcgattctcctgcctcagcctcccgagttgtt<br>gggattccaggcatgcatgaccaggctcagctaatttttgtttt<br>tttggtagagacggggtttcaccatattggccaggctggtctc<br>caactcctaatctcaggtgatctacccaccttggcctcccaa<br>attgctgggattacaggcgtgaaccactgctcccttccctgtc<br>ctt |
| 3’ Untranslated Region | 3’ UTR (mRNA) | The 3’UTR sequence used in IVT mRNA transcripts (not including polyA tail). | CTGGTACTGCATGCACGCAATGCTAGCT<br>GCCCCTTTCCCGTCCTGGGTACCCCGAG<br>TCTCCCCCGACCTCGGGTCCCAGGTATG<br>CTCCCACCTCCACCTGCCCCACTCACCA<br>CCTCTGCTAGTTCCAGACACCTCCCAAG<br>CACGCAGCAATGCAGCTCAAAACGCTTA<br>GCCTAGCCACACCCCCACGGGAAACAG<br>CAGTGATTAACCTTTAGCAATAAACGAAA<br>GTTTAACTAAGCTATACTAACCCCAGGGT<br>TGGTCAATTTCGTGCCAGCCACACCCTG<br>GAGCTAGC |
| ORF | msfGFP[r5M] | Monomeric Superfolder GFP with methionines mutated to other amino acids. Sequence is from start to stop codon of ORF. | atggtgtccaagggcgaggaactgttcaccggcgtggtgc<br>ccatcctggtggaactggacggcgacgtgaacggccaca<br>agttcagcgtgagaggcgagggcgaaggcgacgccaca<br>aacggaaagctgaccctgaagttcatctgcaccaccggca<br>agctgcccgtgccttggcctaccctcgtgaccacactgacct<br>acggcgtgcagtgcttcagcagataccccgaccacatcaa<br>gagacacgatttcttcaagagcgccctgcccgagggctac<br>gtgcaggaacggaccatcagcttcaaggacgacggcacc<br>tacaagacaagagccgaagtgaagttcgagggcgacac<br>cctcgtgaaccggatcgagctgaagggcatcgacttcaaa<br>gaggacggcaacatcctgggccacaagctggagtacaa<br>cttcaacagccacaacgtgtacatcaccgccgacaagca<br>gaaaaacggcatcaaagccaacttcaagatccggcaca<br>acgtggaggacggcagcgtgcagctggccgaccactacc<br>agcagaacacccccatcggagacggccccgtgctgctgc<br>ccgacaaccactacctgagcacacaaagcaagctgagc<br>aaggaccccaacgagaagcgggaccacgccgtgctgct<br>ggaatttgtgaccgccgctggcatcacccacggcaaggac<br>gagctgtacaagtga |
| ORF | msfBFP[r5M] | Monomeric Superfolder BFP with methionines mutated to other amino acids. Sequence is from start to stop codon of ORF. | ATGGTTAGTAAAGGAGAAGAACTCTTCA<br>CCGGCGTAGTTCCGATTCTCGTTGAGCT<br>GGACGGAGACGTTAACGGGCACAAATTC<br>TCTGTAAGGGGTGAAGGGGAAGGCGAC<br>GCTACCAACGGAAAACTGACCTTAAAATT<br>TATTTGTACAACAGGTAAGCTCCCAGTTC<br>CCTGGCCCACCCTTGTTACCACTCTGAC<br>TCACGGCGTTCAGTGTTTCTCTCGTTATC<br>CAGATCATATAAAGCGGCACGACTTCTTT<br>AAGTCCGCTCTCCCCGAAGGCTACGTCC<br>AAGAAAGAACCATTAGTTTTAAGGACGAC<br>GGAACTTACAAAACCAGAGCAGAGGTCA<br>AGTTCGAGGGAGATACTCTCGTCAATCG<br>GATCGAACTGAAAGGAATAGACTTTAAA<br>GAGGACGGCAATATCCTGGGTCACAAGC<br>TGGAGTATAATTTCAATAGCCATAACGTG<br>TATATCACGGCCGATAAGCAGAAGAACG<br>GTATTAAGGCAAATTTTAAGATTCGCCAT<br>AACGTAGAAGACGGTTCCGTCCAATTGG<br>CGGACCATTATCAACAAAACACTCCAATT<br>GGAGACGGCCCCGGTATTACTGCCTGATA<br>ATCACTACCTGTCCACCCAATCAAAACTC<br>TCCAAAGATCCCAACGAGAAACGGGACC |

| | | | |
|---|---|---|---|
| | | | ACGCGGTTCTGTTAGAATTTGTAACGGCCGCTGGTATTACGCACGGTAAGGACGAACTGTATAAGTGA |
| ORF | mEBFP2 | Monomeric enhanced BFP. Sequence is from start to stop codon of ORF. | atggtgtccaagggcgaggaactgttcaccggcgtggtgcccatcctggtggaactggatggcgacgtgaacggccacaagttctctgtgcggggagagggcgaaggcgacgccacaaatggcaagctgaccctgaagttcatctgcaccaccggcaagctgcccgtgccttggcctaccctcgtgaccacactgtctcacggcgtgcagtgcttcgccagataccccgaccacatgaagcagcacgatttcttcaagagcgccatgcccgagggctacgtgcaggaacggaccatcttcttcaaggacgacggcacctacaagaccagagccgaagtgaagttcgagggcgacaccctcgtgaaccggatcgagctgaagggcgtggacttcaaagaggacggcaacatcctgggccacaagctggagtacaacttcaacagccacaacatctacatcatggccgtgaagcagaaaaacggcatcaaagtgaacttcaagatccggcacaacgtggaagatggcagcgtgcagctggccgaccactaccagcagaacacccccatcggagatggccccgtgctgctgcctgatagccactacctgagcacccagagcaagctgagcaaggaccccaacgagaagcgggaccacatggtgctgctggaatttcggaccgccgctggcatcaccctgggcatggatgagctgtacaagtga |
| ORF | emiRFP670 | Far red fluorescent protein with additional +1 Valine mutation to make +1 nucleotide after start codon a guanine nucleotide. Sequence is from start to stop codon of ORF. | atggtggcggaaggctccgtcgccaggcagcctgacctcttgacctgcgaacatgaagagatccacctcgccggctcgatccagccgcatggcgcgcttctggtcgtcagcgaacatgatcatcgcgtcatccaggccagcgccaacgccgcggaatttctgaatctcggaagcgtactcggcgttccgctcgccgagatcgacggcgatctgttgatcaagatcctgccgcatctcgatcccaccgccgaaggcatgccggtcgcggtgcgctgccggatcggcaatccctctacggagtactgcggtctgatgcatcggcctccggaaggcgggctgatcatcgaactcgaacgtgccggcccgtcgatcgatctgtcaggcacgctggcgccggcgctggagcggatccgcacggcgggttcactgcgcgcgctgtgcgatgacaccgtgctgctgtttcagcagtgcaccggctacgaccgggtgatggtgtatcgtttcgatgagcaaggccacggcctggtattctccgagtgccatgtgcctgggctcgaatcctatttcggcaaccgctatccgtcgtcgactgtcccgcagatggcgcggcagctgtacgtgcggcagcgcgtccgcgtgctggtcgacgtcacctatcagccggtgccgctggagccgcggctgtcgccgctgaccgggcgcgatctcgacatgtcgggctgcttcctgcgctcgatgtcgccgtgccatctgcagttcctgaaggacatgggcgtgcgcgccaccctggcggtgtcgctggtggtcggcggcaagctgtggggcctggttgtctgtcaccattatctgccgcgcttcatccgtttcgagctgcgggcgatctgcaaacggctcgccgaaaggatcgcgacgcggatcaccgcgcttgagagctga |
| Internal Ribosome Entry Site | IRES | This IRES sequence was used in all constructs that contained mCherry. | cccccccctaacgttactggccgaagccgcttggaataaggccggtgtgcgtttgtctatatgttattttccaccatattgccgtcttttggcaatgtgagggcccggaaacctggccctgtcttcttgacgagcattcctaggggtctttcccctctcgccaaaggaatgcaaggtctgttgaatgtcgtgaaggaagcagttcctctggaagcttcttgaagacaaacaacgtctgtagcgaccctttgcaggcagcggaaccccccacctggcgacaggtgcctctgcggccaaaagccacgtgtataagatacacctgcaaaggcggcacaaccccagtgccacgttgtgagttggatagttgtggaaagagtcaaatggctctcctcaagcgtattcaacaaggggctgaaggatgcccagaaggtaccccattgtatgggatctgatctggggcctcggtgcacatgctttacatgtgtttagtcgaggttaaaaaacgtctaggccccccgaaccacggggacgtggttttcctttgaaaaacacgatgataatatggccacaacc |

| | | | |
|---|---|---|---|
| Enhancer | CMV Enhancer | Enhancer region to drive expression from CMV promoter. | cgttacataacttacggtaaatggcccgcctggctgaccgc<br>ccaacgacccccgcccattgacgtcaataatgacgtatgtt<br>cccatagtaacgccaatagggactttccattgacgtcaatg<br>ggtggagtatttacggtaaactgcccacttggcagtacatca<br>agtgtatcatatgccaagtacgccccctattgacgtcaatga<br>cggtaaatggcccgcctggcattatgcccagtacatgacctt<br>atgggactttcctacttggcagtacatctacgtattagtcatcg<br>ctattaccatg |
| Promoter | CMV Promoter | CMV Promoter to drive transcription of plasmid-based systems. | gtgatgcggttttggcagtacatcaatgggcgtggatagcg<br>gtttgactcacggggatttccaagtctccaccccattgacgtc<br>aatgggagtttgttttggcaccaaaatcaacgggactttcca<br>aaatgtcgtaacaactccgccccattgacgcaaatgggcg<br>gtaggcgtgtacggtgggaggtctatataagcagagct |
| Promoter | T7 Promoter | T7 Promoter encoded in IVT plasmid backbones. An additional "A" nucleotide is added when PCR amplifying with p101 for capping purposes. | taatacgactcactatagg |
| Full Plasmid | 2-ORF SEMPER TTT/ACC | SEMPER plasmid encoding msfGFP[r5M] and mEBFP2 in TTT/ACC ratio used in transfection. | ttgagatcctttttttctgcgcgtaatctgctgcttgcaaacaaa<br>aaaaccaccgctaccagcggtggtttgtttgccggatcaag<br>agctaccaactctttttccgaaggtaactggcttcagcagag<br>cgcagataccaaatactgttcttctagtgtagccgtagttagg<br>ccaccacttcaagaactctgtagcaccgcctacatacctcg<br>ctctgctaatcctgttaccagtggctgctgccagtggcgataa<br>gtcgtgtcttaccgggttggactcaagacgatagttaccgga<br>taaggcgcagcggtcgggctgaacggggggttcgtgcac<br>acagcccagcttggagcgaacgacctacaccgaactgag<br>atacctacagcgtgagctatgagaaagcgccacgcttccc<br>gaagggagaaaggcggacaggtatccggtaagcggca<br>gggtcggaacaggagagcgcacgagggagcttccaggg<br>ggaaacgcctggtatctttatagtcctgtcgggtttcgccacct<br>ctgacttgagcgtcgatttttgtgatgctcgtcaggggggcgg<br>agcctatggaaaaacgccagcaacgcggcctttttacggtt<br>cctggccttttgctggccttttgctcacatgttctttcctgcgttatt<br>ctttcctgcgttatcccctgattctgtggataaccgtattaccgc<br>ctttgagtgagctgataccgctcgccgcagccgaacgacc<br>gagcgcagcgagtcagtgagcgaggaagcggaagagc<br>gcccaatacgcaaaccgcctctccccgcgcgttggccgatt<br>cattaatgcagagcttgcaattcgcgctttttcaatattattgaa<br>gcatttatcagggttattgtctcatgagcggatacatatttgaa<br>tgtatttagaaaaataaacaaataggggttccgcgcacattt<br>ccccgaaaagtgccacctgacgtctaagaaaccattattat<br>catgacattaacctataaaaataggcgtatcacgaggccct<br>ttcactcattaggcaccccaggctttacactttatgcttccggc<br>tcgtataatgtgtggaattgtgagcggataacaatttcacac<br>aggaaacagcatcgtgcaggtcgttacataacttacggtaa<br>atggcccgcctggctgaccgcccaacgacccccgcccatt<br>gacgtcaataatgacgtatgttcccatagtaacgccaatag<br>ggactttccattgacgtcaatgggtggagtatttacggtaaac<br>tgcccacttggcagtacatcaagtgtatcatatgccaagtac<br>gccccctattgacgtcaatgacggtaaatggcccgcctggc<br>attatgcccagtacatgaccttatgggactttcctacttggca<br>gtacatctacgtattagtcatcgctattaccatggtgatgcggt<br>tttggcagtacatcaatgggcgtggatagcggtttgactcac<br>ggggatttccaagtctccaccccattgacgtcaatgggagtt<br>tgttttggcaccaaaatcaacgggactttccaaaatgtcgta<br>acaactccgccccattgacgcaaatgggcggtaggcgtgt<br>acggtgggaggtctatataagcagagctcgtttagtgaacc<br>gtcagatcgcctggagacgccatccacgctgttttgacctcc<br>atagaagacaccgggaccgatccagcctccggactctag |

| | | | |
|---|---|---|---|
| | | | cctaggcttttgcaaaaagctatttaggtgacactatagaag<br>gtacgcctgcaggtaccgagctcggatccagtagtttaaac<br>cccttctttatggtgtccaagGGCGAGGAACTGTTC<br>ACCGGCGTGGTGCCCATCCTGGTGGAA<br>CTGGACGGCGACGTGAACGGCCACAAG<br>TTCAGCGTGAGAGGCGAGGGCGAAGGC<br>GACGCCACAAACGGAAAGCTGACCCTGA<br>AGTTCATCTGCACCACCGGCAAGCTGCC<br>CGTGCCTTGGCCTACCCTCGTGACCACA<br>CTGACCTACGGCGTGCAGTGCTTCAGCA<br>GATACCCCGACCACATCAAGAGACACGA<br>TTTCTTCAAGAGCGCCCTGCCCGAGGGC<br>TACGTGCAGGAACGGACCATCAGCTTCA<br>AGGACGACGGCACCTACAAGACAAGAG<br>CCGAAGTGAAGTTCGAGGGCGACACCCT<br>CGTGAACCGGATCGAGCTGAAGGGCAT<br>CGACTTCAAAGAGGACGGCAACATCCTG<br>GGCCACAAGCTGGAGTACAACTTCAACA<br>GCCACAACGTGTACATCACCGCCGACAA<br>GCAGAAAAACGGCATCAAAGCCAACTTC<br>AAGATCCGGCACAACGTGGAGGACGGC<br>AGCGTGCAGCTGGCCGACCACTACCAG<br>CAGAACACCCCCATCGGAGACGGCCCC<br>GTGCTGCTGCCCGACAACCACTACCTGA<br>GCACACAAAGCAAGCTGAGCAAGGACCC<br>CAACGAGAAGCGGGACCACGCCGTGCT<br>GCTGGAATTTGTGACCGCCGCTGGCATC<br>ACCCACggcaaggacgagctgtacaagtgacccggg<br>ttaccatggtgtccaagggcgaggaactgttcaccggcgtg<br>gtgcccatcctggtggaactggatggcgacgtgaacggcc<br>acaagttctctgtgcggggagagggcgaaggcgacgcca<br>caaatggcaagctgaccctgaagttcatctgcaccaccgg<br>caagctgcccgtgccttggcctaccctcgtgaccacactgtc<br>tcacggcgtgcagtgcttcgccagataccccgaccacatg<br>aagcagcacgatttcttcaagagcgccatgcccgagggct<br>acgtgcaggaacggaccatcttcttcaaggacgacggcac<br>ctacaagaccagagccgaagtgaagttcgagggcgaca<br>ccctcgtgaaccggatcgagctgaagggcgtggacttcaa<br>agaggacggcaacatcctgggccacaagctggagtaca<br>acttcaacagccacaacatctacatcatggccgtgaagca<br>gaaaaacggcatcaaagtgaacttcaagatccggcacaa<br>cgtggaagatggcagcgtgcagctggccgaccactacca<br>gcagaacacccccatcggagatggccccgtgctgctgcct<br>gatagccactacctgagcacccagagcaagctgagcaag<br>gaccccaacgagaagcgggaccacatggtgctgctgga<br>atttcggaccgccgctggcatcaccctgggcatggatgagc<br>tgtacaagtgataatactaaatcgcactgtcggcgtcccccc<br>ctaacgttactggccgaagccgcttggaataaggccggtgt<br>gcgtttgtctatatgttattttccaccatattgccgtcttttggcaa<br>tgtgagggcccggaaacctggccctgtcttcttgacgagcat<br>tcctaggggtctttcccctctcgccaaaggaatgcaaggtct<br>gttgaatgtcgtgaaggaagcagttcctctggaagcttcttga<br>agacaaacaacgtctgtagcgaccctttgcaggcagcgg<br>aacccccacctggcgacaggtgcctctgcggccaaaag<br>ccacgtgtataagatacacctgcaaaggcggcacaaccc<br>cagtgccacgttgtgagttggatagttgtggaaagagtcaa<br>atggctctcctcaagcgtattcaacaaggggctgaaggatg<br>cccagaaggtaccccattgtatgggatctgatctggggcctc<br>ggtgcacatgctttacatgtgtttagtcgaggttaaaaaacgt<br>ctaggccccccgaaccacggggacgtggttttcctttgaaa<br>aacacgatgataatatggccacaaccatggtgagcaagg<br>gcgaggaggataacatggccatcatcaaggagttcatgcg<br>cttcaaggtgcacatggagggctccgtgaacggccacga<br>gttcgagatcgagggcgagggcgagggccgcccctacg<br>agggcacccagaccgccaagctgaaggtgaccaagggc |

| | | | |
|---|---|---|---|
| | | | ggccccctgcccttcgcctgggacatcctgtcccctcagttc<br>atgtacggctccaaggcctacgtgaagcaccccgccgac<br>atccccgactacttgaagctgtccttccccgagggcttcaag<br>tgggagcgcgtgatgaacttcgaggacggcggcgtggtga<br>ccgtgacccaggactcctccctgcaggacggcgagttcat<br>ctacaaggtgaagctgcgcggcaccaacttcccctccgac<br>ggccccgtaatgcagaagaagaccatgggctgggaggc<br>ctcctccgagcggatgtaccccgaggacggcgccctgaa<br>gggcgagatcaagcagaggctgaagctgaaggacggcg<br>gccactacgacgctgaggtcaagaccacctacaaggcca<br>agaagcccgtgcagctgcccggcgcctacaacgtcaaca<br>tcaagttggacatcacctcccacaacgaggactacaccat<br>cgtggaacagtacgaacgcgccgagggccgccactcca<br>ccggcggcatggatgagctgtacaagtgaaggatccctcg<br>aggggcccaagcttacgcgtgcatgcgacgtcatagctctc<br>tccctatagtgagtcgtattataagctagcttgggatctttgtga<br>aggaaccttacttctgtggtgtgacataattggacaaactac<br>ctacagagatttaaagctctaaggtaaatataaaatttttaag<br>tgtataatgtgttaaactagctgcatatgcttgctgcttgagag<br>ttttgcttactgagtatgatttatgaaaatattatacacaggag<br>ctagtgattctaattgtttgtgtattttagattcacagtcccaagg<br>ctcatttcaggcccctcagtcctcacagtctgttcatgatcata<br>atcagccataccacatttgtagaggttttacttgctttaaaaaa<br>cctcccacacctccccctgaacctgaaacataaaatgaat<br>gcaattgttgttgttcgtggctgaatcaacctctggattacaaa<br>atttgtgaaagattgactggtattcttaactatgttgctccttttac<br>gctatgtggatacgctgctttaatgcctttgtatcatgctattgct<br>tcccgtatggctttcattttctcctccttgtataaatcctggttgct<br>gtctctttatgaggagttgtggcccgttgtcaggcaacgtggc<br>gtggtgtgcactgtgtttgctgacgcaacccccactggttgg<br>ggcattgccaccacctgtcagctcctttccgggactttcgcttt<br>ccccctccctattgccacggcggaactcatcgccgcctgcc<br>ttgcccgctgctggacaggggctcggctgttgggcactgac<br>aattccgtggtgttgtcggggaaatcatcgtcctttccttggct<br>gctcgcctgtgttgccacctggattctgcgcgggacgtccttc<br>tgctacgtcccttcggccctcaatccagcggaccttccttccc<br>gcggcctgctgccggctctgcggcctcttccgcgtcttcgcct<br>tcgccctcagacgagtcggatctccctttgggccgcctcccc<br>gccagagacaattaacttcgcgggtggcatccctgtgaccc<br>ctccccagtgcctctcctggccctggaagttgccactccagt<br>gcccaccagccttgtcctaataaaattaagttgcatcattttgt<br>ctgactaggtgtccttctataatattatggggtggaggggggt<br>ggtatggagcaaggggcaagttgggaagacaacctgtag<br>ggcctgcggggtctattgggaaccaagctggagtgcagtg<br>gcacaatcttggctcactgcaatctccgcctcctgggttcaa<br>gcgattctcctgcctcagcctcccgagttgttgggattccagg<br>catgcatgaccaggctcagctaatttttgtttttttggtagagac<br>ggggtttcaccatattggccaggctggtctccaactcctaatc<br>tcaggtgatctacccaccttggcctcccaaattgctgggatta<br>caggcgtgaaccactgctcccttccctgtccttgaagttctca<br>gatcctgcattaatgaatcggccaacgcgcggggagagg<br>cggtttgcgtattggctggcgtaatagcgaagaggcccgca<br>ccgatcgcccttcccaacagttgcgcagcctgaatggcga<br>atgggacgcgccctgtagcggcgcattaagcgcggcggg<br>tgtggtggttacgcgcagcgtgaccgctacacttgccagcg<br>ccctagcgcccgctcctttcgctttcttcccttcctttctcgccac<br>gttcgccggctttccccgtcaagctctaaatcgggggctccc<br>tttagggttccgatttagtgctttacggcacctcgaccccaaa<br>aaacttgattagggtgatggttcacgtagtgggccatcgccc<br>tgatagacggtttttcgccctttgacgttggagtccacgttctta<br>atagtggactcttgttccaaactggaacaacactcaaccta<br>tctcggtctattcttttgatttataagggattttgccgatttcggcc<br>tattggttaaaaaatgagctgatttaacaaaaatttaacgcg<br>aattttaacaaaatattaacgcttacaatttaggtggcacttttc<br>ggggaaatgtgcgcggaacccctatttgtttatttttctaaata |

| | | | |
|---|---|---|---|
| | | | cattcaaatatgtatccgctcatgccaggtcttggactggtga<br>gaacggcttgctcggcagcttcgatgtgtgctggagggaga<br>ataaaggtctaagatgtgcgatagagggaagtcgcattga<br>attatgtgctgtgtagggatcgctggtatcaaatatgtgtgccc<br>acccctggcatgagacaataaccctgataaatgcttcaata<br>atattgaaaaaggaagagtatgagtattcaacatttccgtgt<br>cgcccttattcccttttttgcggcattttgccttcctgtttttgctca<br>cccagaaacgctggtgaaagtaaaagatgctgaagatca<br>gttgggtgcacgagtgggttacatcgaactggatctcaaca<br>gcggtaagatccttgagagttttcgccccgaagaacgttttc<br>caatgatgagcacttttaaagttctgctatgtggcgcggtatta<br>tcccgtattgacgccgggcaagagcaactcggtcgccgca<br>tacactattctcagaatgacttggttgagtactcaccagtcac<br>agaaaagcatcttacggatggcatgacagtaagagaatta<br>tgcagtgctgccataaccatgagtgataacactgcggcca<br>acttacttctgacaacgatcggaggaccgaaggagctaac<br>cgcttttttgcacaacatgggggatcatgtaactcgccttgat<br>cgttgggaaccggagctgaatgaagccataccaaacgac<br>gagcgtgacaccacgatgcctgtagcaatggcaacaacg<br>ttgcgcaaactattaactggcgaactacttactctagcttccc<br>ggcaacaattaatagactggatggaggcggataaagttgc<br>aggaccacttctgcgctcggcccttccggctggctggtttatt<br>gctgataaatctggagccggtgagcgtgggtctcgcggtat<br>cattgcagtactggggccagatggtaagccctcccgtatcgt<br>agttatctacacgacggggagtcaggcaactatggatgaa<br>cgaaatagacagatcgctgagataggtgcctcactgattaa<br>gcattggtaactgtcagaccaagtttactcatatatactttaga<br>ttgatttaaaacttcatttttaatttaaaaggatctaggtgaaga<br>tcctttttgataatctcatgccataacttcgtataatgtatgctat<br>acgaagttatggcatgaccaaaatcccttaacgtgagttttc<br>gttccactgagcgtcagaccccgtagaaaagatcaaagg<br>atcttc |
| Full Plasmid | 3-ORF SEMPER Plasmid ACC/***/*** | 3-ORF SEMPER plasmid used in transient transfection experiments. | ttgagatcctttttttctgcgcgtaatctgctgcttgcaaacaaa<br>aaaaccaccgctaccagcggtggtttgtttgccggatcaag<br>agctaccaactctttttccgaaggtaactggcttcagcagag<br>cgcagataccaaatactgttcttctagtgtagccgtagttagg<br>ccaccacttcaagaactctgtagcaccgcctacatacctcg<br>ctctgctaatcctgttaccagtggctgctgccagtggcgataa<br>gtcgtgtcttaccgggttggactcaagacgatagttaccgga<br>taaggcgcagcggtcgggctgaacggggggttcgtgcac<br>acagcccagcttggagcgaacgacctacaccgaactgag<br>atacctacagcgtgagctatgagaaagcgccacgcttccc<br>gaagggagaaaggcggacaggtatccggtaagcggca<br>gggtcggaacaggagagcgcacgagggagcttccaggg<br>ggaaacgcctggtatctttatagtcctgtcgggtttcgccacct<br>ctgacttgagcgtcgatttttgtgatgctcgtcaggggggcgg<br>agcctatggaaaaacgccagcaacgcggcctttttacggtt<br>cctggccttttgctggccttttgctcacatgttctttcctgcgttatt<br>ctttcctgcgttatcccctgattctgtggataaccgtattaccgc<br>ctttgagtgagctgataccgctcgccgcagccgaacgacc<br>gagcgcagcgagtcagtgagcgaggaagcggaagagc<br>gcccaatacgcaaaccgcctctccccgcgcgttggccgatt<br>cattaatgcagagcttgcaattcgcgctttttcaatattattgaa<br>gcatttatcagggttattgtctcatgagcggatacatatttgaa<br>tgtatttagaaaaataaacaaataggggttccgcgcacattt<br>ccccgaaaagtgccacctgacgtctaagaaaccattattat<br>catgacattaacctataaaaataggcgtatcacgaggccct<br>ttcactcattaggcaccccaggctttacactttatgcttccggc<br>tcgtataatgtgtggaattgtgagcggataacaatttcacac<br>aggaaacagcatcgtgcaggtcgttacataacttacggtaa<br>atggcccgcctggctgaccgcccaacgacccccgcccatt<br>gacgtcaataatgacgtatgttcccatagtaacgccaatag<br>ggactttccattgacgtcaatgggtggagtatttacggtaaac |

| | | | |
|---|---|---|---|
| | | | tgcccacttggcagtacatcaagtgtatcatatgccaagtac<br>gccccctattgacgtcaatgacggtaaatggcccgcctggc<br>attatgcccagtacatgaccttatgggactttcctacttggca<br>gtacatctacgtattagtcatcgctattaccatggtgatgcggt<br>tttggcagtacatcaatgggcgtggatagcggtttgactcac<br>ggggatttccaagtctccaccccattgacgtcaatgggagtt<br>tgttttggcaccaaaatcaacgggactttccaaaatgtcgta<br>acaactccgccccattgacgcaaatgggcggtaggcgtgt<br>acggtgggaggtctatataagcagagctcgtttagtgaacc<br>gtcagatcgcctggagacgccatccacgctgttttgacctcc<br>atagaagacaccgggaccgatccagcctccggactctag<br>cctaggcttttgcaaaaagctatttaggtgacactatagaag<br>gtacgcctgcaggtaccgagctcggatccagtagtttaaac<br>cccttcACCATGGTTAGTAAAGGAGAAGAA<br>CTCTTCACCGGCGTAGTTCCGATTCTCG<br>TTGAGCTGGACGGAGACGTTAACGGGCA<br>CAAATTCTCTGTAAGGGGTGAAGGGGAA<br>GGCGACGCTACCAACGGAAAACTGACCT<br>TAAAATTTATTTGTACAACAGGTAAGCTC<br>CCAGTTCCCTGGCCCACCCTTGTTACCA<br>CTCTGACTCACGGCGTTCAGTGTTTCTCT<br>CGTTATCCAGATCATATAAAGCGGCACG<br>ACTTCTTTAAGTCCGCTCTCCCCGAAGG<br>CTACGTCCAAGAAAGAACCATTAGTTTTA<br>AGGACGACGGAACTTACAAAACCAGAGC<br>AGAGGTCAAGTTCGAGGGAGATACTCTC<br>GTCAATCGGATCGAACTGAAAGGAATAG<br>ACTTTAAAGAGGACGGCAATATCCTGGG<br>TCACAAGCTGGAGTATAATTTCAATAGCC<br>ATAACGTGTATATCACGGCCGATAAGCA<br>GAAGAACGGTATTAAGGCAAATTTTAAGA<br>TTCGCCATAACGTAGAAGACGGTTCCGT<br>CCAATTGGCGGACCATTATCAACAAAAC<br>ACTCCAATTGGAGACGGCCCGGTATTAC<br>TGCCTGATAATCACTACCTGTCCACCCAA<br>TCAAAACTCTCCAAAGATCCCAACGAGA<br>AACGGGACCACGCGGTTCTGTTAGAATT<br>TGTAACGGCCGCTGGTATTACGCACGGT<br>AAGGACGAACTGTATAAGTGAAGCGCTtt<br>TTTCCATtgTCCAAGGGCGAGGAACTGTT<br>CACCGGCGTGGTGCCCATCCTGGTGGA<br>ACTGGACGGCGACGTGAACGGCCACAA<br>GTTCAGCGTGAGAGGCGAGGGCGAAGG<br>CGACGCCACAAACGGAAAGCTGACCCTG<br>AAGTTCATCTGCACCACCGGCAAGCTGC<br>CCGTGCCTTGGCCTACCCTCGTGACCAC<br>ACTGACCTACGGCGTGCAGTGCTTCAGC<br>AGATACCCCGACCACATCAAGAGACACG<br>ATTTCTTCAAGAGCGCCCTGCCCGAGGG<br>CTACGTGCAGGAACGGACCATCAGCTTC<br>AAGGACGACGGCACCTACAAGACAAGAG<br>CCGAAGTGAAGTTCGAGGGCGACACCCT<br>CGTGAACCGGATCGAGCTGAAGGGCAT<br>CGACTTCAAAGAGGACGGCAACATCCTG<br>GGCCACAAGCTGGAGTACAACTTCAACA<br>GCCACAACGTGTACATCACCGCCGACAA<br>GCAGAAAAACGGCATCAAAGCCAACTTC<br>AAGATCCGGCACAACGTGGAGGACGGC<br>AGCGTGCAGCTGGCCGACCACTACCAG<br>CAGAACACCCCCATCGGAGACGGCCCC<br>GTGCTGCTGCCCGACAACCACTACCTGA<br>GCACACAAAGCAAGCTGAGCAAGGACCC<br>CAACGAGAAGCGGGACCACGCCGTGCT<br>GCTGGAATTTGTGACCGCCGCTGGCATC<br>ACCCACGgcaaggacgagctgtacaagtgacccggg |

| | | | |
|---|---|---|---|
| | | | ttTTTCCATtggcggaaggctccgtcgccaggcagcct<br>gacctcttgacctgcgaacatgaagagatccacctcgccg<br>gctcgatccagccgcatggcgcgcttctggtcgtcagcgaa<br>catgatcatcgcgtcatccaggccagcgccaacgccgcg<br>gaatttctgaatctcggaagcgtactcggcgttccgctcgcc<br>gagatcgacggcgatctgttgatcaagatcctgccgcatctc<br>gatcccaccgccgaaggcatgccggtcgcggtgcgctgc<br>cggatcggcaatccctctacggagtactgcggtctgatgcat<br>cggcctccggaaggcgggctgatcatcgaactcgaacgtg<br>ccggcccgtcgatcgatctgtcaggcacgctggcgccggc<br>gctggagcggatccgcacggcgggttcactgcgcgcgctg<br>tgcgatgacaccgtgctgctgtttcagcagtgcaccggctac<br>gaccgggtgatggtgtatcgtttcgatgagcaaggccacgg<br>cctggtattctccgagtgccatgtgcctgggctcgaatcctatt<br>tcggcaaccgctatccgtcgtcgactgtcccgcagatggcg<br>cggcagctgtacgtgcggcagcgcgtccgcgtgctggtcg<br>acgtcacctatcagccggtgccgctggagccgcggctgtc<br>gccgctgaccgggcgcgatctcgacatgtcgggctgcttcc<br>tgcgctcgatgtcgccgtgccatctgcagttcctgaaggaca<br>tgggcgtgcgcgccaccctggcggtgtcgctggtggtcggc<br>ggcaagctgtggggcctggttgtctgtcaccattatctgccgc<br>gcttcatccgtttcgagctgcgggcgatctgcaaacggctcg<br>ccgaaaggatcgcgacgcggatcaccgcgcttgagagct<br>gataaTCTAGAtacactaaatcgcactgtcggcgtcccc<br>ccctaacgttactggccgaagccgcttggaataaggccggt<br>gtgcgtttgtctatatgttattttccaccatattgccgtcttttggc<br>aatgtgagggcccggaaacctggccctgtcttcttgacgag<br>cattcctaggggtctttcccctctcgccaaaggaatgcaagg<br>tctgttgaatgtcgtgaaggaagcagttcctctggaagcttctt<br>gaagacaaacaacgtctgtagcgaccctttgcaggcagc<br>ggaaccccccacctggcgacaggtgcctctgcggccaaa<br>agccacgtgtataagatacacctgcaaaggcggcacaac<br>cccagtgccacgttgtgagttggatagttgtggaaagagtca<br>aatggctctcctcaagcgtattcaacaaggggctgaaggat<br>gcccagaaggtaccccattgtatgggatctgatctggggcc<br>tcggtgcacatgctttacatgtgtttagtcgaggttaaaaaac<br>gtctaggccccccgaaccacggggacgtggttttcctttgaa<br>aaacacgatgataatatggccacaaccatggtgagcaag<br>ggcgaggaggataacatggccatcatcaaggagttcatgc<br>gcttcaaggtgcacatggagggctccgtgaacggccacg<br>agttcgagatcgagggcgagggcgagggccgcccctac<br>gagggcacccagaccgccaagctgaaggtgaccaagg<br>gcggcccccctgcccttcgcctgggacatcctgtcccctcagt<br>tcatgtacggctccaaggcctacgtgaagcaccccgccga<br>catccccgactacttgaagctgtccttccccgagggcttcaa<br>gtgggagcgcgtgatgaacttcgaggacggcggcgtggtg<br>accgtgacccaggactcctccctgcaggacggcgagttca<br>tctacaaggtgaagctgcgcggcaccaacttcccctccga<br>cggccccgtaatgcagaagaagaccatgggctgggagg<br>cctcctccgagcggatgtaccccgaggacggcgccctga<br>agggcgagatcaagcagaggctgaagctgaaggacggc<br>ggccactacgacgctgaggtcaagaccacctacaaggcc<br>aagaagcccgtgcagctgcccggcgcctacaacgtcaac<br>atcaagttggacatcacctcccacaacgaggactacacca<br>tcgtggaacagtacgaacgcgccgagggccgccactcca<br>ccggcggcatggatgagctgtacaagtgaaggatccctcg<br>aggggcccaagcttacgcgtgcatgcgacgtcatagctctc<br>tccctatagtgagtcgtattataagctagcttgggatctttgtga<br>aggaaccttacttctgtggtgtgacataattggacaaactac<br>ctacagagatttaaagctctaaggtaaatataaaatttttaag<br>tgtataatgtgttaaactagctgcatatgcttgctgcttgagag<br>ttttgcttactgagtatgatttatgaaaatattatacacaggag<br>ctagtgattctaattgtttgtgtattttagattcacagtcccaagg<br>ctcatttcaggcccctcagtcctcacagtctgttcatgatcata<br>atcagccataccacatttgtagaggttttacttgctttaaaaaa |

| | | | |
|---|---|---|---|
| | | | cctcccacacctccccctgaacctgaaacataaaatgaat<br>gcaattgttgttgttcgtggctgaatcaacctctggattacaaa<br>atttgtgaaagattgactggtattcttaactatgttgctccttttac<br>gctatgtggatacgctgctttaatgcctttgtatcatgctattgct<br>tcccgtatggctttcattttctcctccttgtataaatcctggttgct<br>gtctctttatgaggagttgtggcccgttgtcaggcaacgtggc<br>gtggtgtgcactgtgtttgctgacgcaacccccactggttgg<br>ggcattgccaccacctgtcagctcctttccgggactttcgcttt<br>ccccctccctattgccacggcggaactcatcgccgcctgcc<br>ttgcccgctgctggacaggggctcggctgttgggcactgac<br>aattccgtggtgttgtcggggaaatcatcgtcctttccttggct<br>gctcgcctgtgttgccacctggattctgcgcgggacgtccttc<br>tgctacgtcccttcggccctcaatccagcggaccttccttccc<br>gcggcctgctgccggctctgcggcctcttccgcgtcttcgcct<br>tcgccctcagacgagtcggatctccctttgggccgcctcccc<br>gccagagacaattaacttcgcgggtggcatccctgtgaccc<br>ctccccagtgcctctcctggccctggaagttgccactccagt<br>gcccaccagccttgtcctaataaaattaagttgcatcattttgt<br>ctgactaggtgtccttctataatattatggggtggaggggggt<br>ggtatggagcaaggggcaagttgggaagacaacctgtag<br>ggcctgcggggtctattgggaaccaagctggagtgcagtg<br>gcacaatcttggctcactgcaatctccgcctcctgggttcaa<br>gcgattctcctgcctcagcctcccgagttgttgggattccagg<br>catgcatgaccaggctcagctaatttttgtttttttggtagagac<br>ggggtttcaccatattggccaggctggtctccaactcctaatc<br>tcaggtgatctacccaccttggcctcccaaattgctgggatta<br>caggcgtgaaccactgctcccttccctgtccttgaagttctca<br>gatcctgcattaatgaatcggccaacgcgcggggagagg<br>cggtttgcgtattggctggcgtaatagcgaagaggcccgca<br>ccgatcgcccttcccaacagttgcgcagcctgaatggcga<br>atgggacgcgccctgtagcggcgcattaagcgcggcggg<br>tgtggtggttacgcgcagcgtgaccgctacacttgccagcg<br>ccctagcgcccgctcctttcgctttcttcccttcctttctcgccac<br>gttcgccggctttccccgtcaagctctaaatcgggggctccc<br>tttagggttccgatttagtgctttacggcacctcgaccccaaa<br>aaacttgattagggtgatggttcacgtagtgggccatcgccc<br>tgatagacggtttttcgccctttgacgttggagtccacgttctta<br>atagtggactcttgttccaaactggaacaacactcaacccta<br>tctcggtctattcttttgatttataagggattttgccgatttcggcc<br>tattggttaaaaaatgagctgatttaacaaaaatttaacgcg<br>aattttaacaaaatattaacgcttacaatttaggtggcacttttc<br>ggggaaatgtgcgcggaacccctatttgtttatttttctaaata<br>cattcaaatatgtatccgctcatgccaggtcttggactggtga<br>gaacggcttgctcggcagcttcgatgtgtgctggagggaga<br>ataaaggtctaagatgtgcgatagagggaagtcgcattga<br>attatgtgctgtgtagggatcgctggtatcaaatatgtgtgccc<br>acccctggcatgagacaataaccctgataaatgcttcaata<br>atattgaaaaaggaagagtatgagtattcaacatttccgtgt<br>cgcccttattcccttttttgcggcattttgccttcctgtttttgctca<br>cccagaaacgctggtgaaagtaaaagatgctgaagatca<br>gttgggtgcacgagtgggttacatcgaactggatctcaaca<br>gcggtaagatccttgagagttttcgccccgaagaacgttttc<br>caatgatgagcacttttaaagttctgctatgtggcgcggtatta<br>tcccgtattgacgccgggcaagagcaactcggtcgccgca<br>tacactattctcagaatgacttggttgagtactcaccagtcac<br>agaaaagcatcttacggatggcatgacagtaagagaatta<br>tgcagtgctgccataaccatgagtgataacactgcggcca<br>acttacttctgacaacgatcggaggaccgaaggagctaac<br>cgcttttttgcacaacatgggggatcatgtaactcgccttgat<br>cgttgggaaccggagctgaatgaagccataccaaacgac<br>gagcgtgacaccacgatgcctgtagcaatggcaacaacg<br>ttgcgcaaactattaactggcgaactacttactctagcttccc<br>ggcaacaattaatagactggatggaggcggataaagttgc<br>aggaccacttctgcgctcggcccttccggctggctggtttatt<br>gctgataaatctggagccggtgagcgtgggtctcgcggtat |

| | | | |
|---|---|---|---|
| | | | cattgcagtactggggccagatggtaagccctcccgtatcgt<br>agttatctacacgacggggagtcaggcaactatggatgaa<br>cgaaatagacagatcgctgagataggtgcctcactgattaa<br>gcattggtaactgtcagaccaagtttactcatatatactttaga<br>ttgatttaaaacttcatttttaatttaaaaggatctaggtgaaga<br>tcctttttgataatctcatgccataacttcgtataatgtatgctat<br>acgaagttatggcatgaccaaaatcccttaacgtgagttttc<br>gttccactgagcgtcagaccccgtagaaaagatcaaagg<br>atcttc |
| Full Plasmid | SEMPER mARG | Full sequence for SEMPER mARG plasmid for the ACC/ACC TIS combination. | ttgagatcctttttttctgcgcgtaatctgctgcttgcaaacaaa<br>aaaaccaccgctaccagcggtggtttgtttgccggatcaag<br>agctaccaactctttttccgaaggtaactggcttcagcagag<br>cgcagataccaaatactgttcttctagtgtagccgtagttagg<br>ccaccacttcaagaactctgtagcaccgcctacatacctcg<br>ctctgctaatcctgttaccagtggctgctgccagtggcgataa<br>gtcgtgtcttaccgggttggactcaagacgatagttaccgga<br>taaggcgcagcggtcgggctgaacggggggttcgtgcac<br>acagcccagcttggagcgaacgacctacaccgaactgag<br>atacctacagcgtgagctatgagaaagcgccacgcttccc<br>gaagggagaaaggcggacaggtatccggtaagcggca<br>gggtcggaacaggagagcgcacgagggagcttccaggg<br>ggaaacgcctggtatctttatagtcctgtcgggtttcgccacct<br>ctgacttgagcgtcgatttttgtgatgctcgtcaggggggcgg<br>agcctatggaaaaacgccagcaacgcggcctttttacggtt<br>cctggccttttgctggccttttgctcacatgttctttcctgcgttatt<br>ctttcctgcgttatcccctgattctgtggataaccgtattaccgc<br>ctttgagtgagctgataccgctcgccgcagccgaacgacc<br>gagcgcagcgagtcagtgagcgaggaagcggaagagc<br>gcccaatacgcaaaccgcctctccccgcgcgttggccgatt<br>cattaatgcagagcttgcaattcgcgctttttcaatattattgaa<br>gcatttatcagggttattgtctcatgagcggatacatatttgaa<br>tgtatttagaaaaataaacaaataggggttccgcgcacattt<br>ccccgaaaagtgccacctgacgtctaagaaaccattattat<br>catgacattaacctataaaaataggcgtatcacgaggccct<br>ttcactcattaggcaccccaggctttacactttatgcttccggc<br>tcgtataatgtgtggaattgtgagcggataacaatttcacac<br>aggaaacagcatcgtgcaggtcgttacataacttacggtaa<br>atggcccgcctggctgaccgcccaacgacccccgcccatt<br>gacgtcaataatgacgtatgttcccatagtaacgccaatag<br>ggactttccattgacgtcaatgggtggagtatttacggtaaac<br>tgcccacttggcagtacatcaagtgtatcatatgccaagtac<br>gccccctattgacgtcaatgacggtaaatggcccgcctggc<br>attatgcccagtacatgaccttatgggactttcctacttggca<br>gtacatctacgtattagtcatcgctattaccatggtgatgcggt<br>tttggcagtacatcaatgggcgtggatagcggtttgactcac<br>ggggatttccaagtctccaccccattgacgtcaatgggagtt<br>tgttttggcaccaaaatcaacgggactttccaaaatgtcgta<br>acaactccgccccattgacgcaaatgggcggtaggcgtgt<br>acggtgggaggtctatataagcagagctcgtttagtgaacc<br>gtcagatcgcctggagacgccatccacgctgttttgacctcc<br>atagaagaccgagctcggatccagtacccttcaccatggc<br>cgtggaaaagaccaacagcagcagctccctggccgaagt<br>gatcgacagaatcctggacaagggcatcgtgatcgacgc<br>ctgggtgcgcgtgtccctcgtgggaattgagctgctggccat<br>cgaggcccggatcgtgattgccagcgtggaaacctacctg<br>aagtacgccgaggccgtgggcctgacacagagtgctgct<br>gtgcctgcttgattaccatggtgaccaccaccaaagtgaac<br>cacaagcgggccgtgctgagactgaggcctggccagtttg<br>tcgtgacccccgccattgagcgggtggccattagagccctg<br>agatacctgaagtccggcttccccgtgcacctgagaggac<br>ctgccggaaccggcaagaccacactggccatgcacctgg<br>ccaactgcctggacagacccgtgatgctgctgttcggcgac<br>gaccagttcaagagcagcgacctgatcggcagcgagag |

| | | | |
|---|---|---|---|
| | | | cggctacacccacaagaaggtgctggacaactacatcca<br>cagcgtcgtgaagctggaagatgagttcaagcagaactgg<br>gtggacagcagactgaccctggcctgccgggaaggcttca<br>ccctggtgtacgacgagttcaaccggtccagacccgaagt<br>gaacaacgtgctgctgagcgccctggaagagaagatcct<br>gagcctgccccccagcagcaaccagccagagtacctga<br>gcgtgaacccccagttcagagtgatcttcaccagcaaccc<br>cgaggaatatgccggcgtgcacagcacccaggacgccct<br>gatggaccggctcgtgaccatctccatgcccgagcccgat<br>gagatcacccagaccgagatcctgatccagaaaacaaac<br>atcgaccgcgagagcgccaacttcatcgtgcggctcgtga<br>agtccttcagactggccacaggcgccgagaaaaccagcg<br>gcctgagaagctgcctgatgatcgccaaagtgtgcgccga<br>caacaacatccccgtgaccaccgagagcctggacttcccc<br>gatatcgccatcgacatcctgttcaaccgcagccacctgag<br>catgagcgagtccaccaacatctttctggaactgctggata<br>agttcagcgccgaggaactggaaatcctgaacaacagag<br>tgaccggcgacaacgacttcctgatcgacaacagccagtt<br>cgtgtcccagcagctggccggacagcccaacggagcgc<br>caggttccggggctactaacttcagcctccttaaacaggctg<br>gcgatgtggaagaaaatcccggaccagtgctgcccacca<br>gaccccagaccaacagcagccggaccatcaacaccagc<br>acccagggcagcaccctggccgacatcctggaaagagtg<br>ctggacaagggcatcgtgatcgccggggacatcagcatct<br>ctatcgccagcaccgagctggtgcacatccggatcagact<br>gctgatctccagcgtggacaaggccaaagagatgggcat<br>caactggtgggagagcgacccctacctgagcaccaaggc<br>ccagagactggtggaagagaaccagcagctccagcaca<br>gactggaaagcctggaagccaagctgaacagcctgacc<br>agcagcagcgtgaaagaggaaatcccccctggccgccga<br>cgtgaaggacgacctgtatcagaccagcgccaagatccc<br>cagccccgtggataccccctatcgaggtgctggacttccagg<br>cccagtctagcggcggcacacccccttacgtgaacacctct<br>atggaaatactggattttcaggctcagacctccgccgagag<br>cagcagccctgtgggctctaccgtggaaatactggacttcc<br>aggcacagacaagcgaggaatccagctcccccgtggtgt<br>ccacagtggaaatactggatttccaggcccagacttctgaa<br>gagtccagctctccagtgggaagcactgtggaaatactgg<br>actttcaggcccagaccagtgaagagatccccagctctgtg<br>gaccccgccatcgatgtgggtgccccgggatctggcgcaa<br>caaattttagtctcctcaagcaggcaggagatgtcgaggaa<br>aaccctggacccgtggtgtgcacccctgccgagaacttca<br>acaacagcctgacaatcgccagcaagcccaagaacgag<br>gccggactggctcctctgctgctgaccgtgctggaactcgtg<br>cggcagctgatggaagcccaagtgatccggcggatggaa<br>gaggacctgctgagcgagcccgacctggaaagagccgc<br>cgatagcctccagaagctggaagaacagatcctgcacctg<br>tgcgagatgttcgaggtggaccccgccgacctgaacatca<br>acctgggcgagatcggcaccctgctgcctagcagcggca<br>gctactatcctggccagccaagcagcagaccctctgtgctg<br>gaactgctggaccggctgctgaacaccggcatcgtggtgg<br>acggcgagattgacctgggaatcgcccagatcgacctgat<br>ccacgccaagctgagactggtgctgacctccaagcccatc<br>ggggcacctggctcaggagcgaccaacttctcattgctcaa<br>acaagccggagatgttgaggagaatccaggccctgtgag<br>catccccctgtacctgtacggcatcttccccaacaccatccc<br>cgagacactggaactggaaggcctggacaagcagcccg<br>tgcacagccaggtggtggacgagttctgcttcctgtacagc<br>gaggcccggcaggaaaagtacctggccagcagacgga<br>acctgctgacccacgagaaggtgctggaacagaccatgc<br>acgccggcttcagagtgctgctgcccctgagatttggcctgg<br>tcgtgaaggactgggagacaatcatgagccagctgatcaa<br>cccccacaaggaccagctgaaccagctgttccagaagct<br>ggccggcaagcgggaagtgtccatcaagattttctgggac<br>gccaaggccgagcttcagaccatgatggaaagccaccag |

| | | | |
|---|---|---|---|
| | | | gacctgaagcagcagcgggacaacatggaaggcaaga<br>aactgagcatggaagaagtgatccagatcgggcagctgat<br>cgagatcaacctgctggcccggaagcaggccgtgatcga<br>ggtgttcagccaggaactgaaccccttcgcccaggaaatc<br>gtggtgtccgaccccatgaccgaggaaatgatctacaacg<br>ccgccttcctgatcccctgggagagcgagagcgagttcag<br>cgagcgggtggaagtgatcgaccagaagttcggcgaccg<br>gctgcggatcagatacaacaacttcaccgccccctacacct<br>ttgcccagctggactctggcgctcccgggtcaggtgccact<br>aatttttcattgttgaagcaagctggggatgttgaagagaac<br>ccagggcctgtgctgaccaaactcctgctgctgcccatcat<br>gggacccctgaatggcgtcgtgtggatcgccgagcagatc<br>caggaacggaccaacaccgagttcgacgcccaggaaaa<br>cctgcacaaacagctgctgagcctccagctgagcttcgac<br>atcggcgagatcggagaggaagagttcgagattcaggaa<br>gaggaaatcctgctgaagatccaggccctggaagaagag<br>gcccggctggaactggaagccgagcaggaagaagcca<br>gactggaactggaagctgaacaggaagatttcgagtaccc<br>accacagttcaccgccgaagtgaacaaggatcagcatctg<br>gtgctgctgcccggtgccccgggatctggcgctactaacttc<br>agcctccttaaacaggctggcgatgtggaagaaaatcccg<br>gaccagtggaactggaaaacctgtacacctacgcctttctg<br>gaaatccccagcagcccccctgatcctgcctcagggcgctg<br>ctaatcaggtggtgctgatcaacggcaccgagctggccgc<br>catcgtggaacctggcatcttcctggaatccttccagaacaa<br>cgacgagaagatcatccagatggccctgagccacgacag<br>agtgatctgcgagctgttccagcagatcaccgtgctgcccct<br>gagattcggcacctacttcaccagcaccaacaacctgctg<br>aaccacctgaagtcccacgagaaagagtaccagaacaa<br>gctggaaaagatcaacgggaagaacgagttcaccctgaa<br>gctgatcccccggatgatcgaggaaatcgtgccctctgagg<br>gcggaggcaaggactacttcctggccaagaagcagcgct<br>accagaatcagaacaacttctctatcgcccaggccgccga<br>gaagcagaacctgatcgacctgatcaccaaagtgaacca<br>gctgcccgtggtggtgcaggaacaggaagaacagatcca<br>aatctacctgctggtgtcctgccaggataagaccctgctgct<br>ggaacagttcctgacctggcagaaagcctgccccaagtgg<br>gatctgctgctgggcgactgcctgccccccctaccactttatcg<br>gagcgccaggttccggggcaacaaattttagtctcctcaag<br>caggcaggagatgtcgaggaaaaccctggacccgtggtg<br>tccaagggcgaggaactgttcaccggcgtggtgcccatcct<br>ggtggaactggatggcgacgtgaacggccacaagttcag<br>cgtgtccggcgagggcgaaggcgacgccacatacggaa<br>agctgaccctgaagttcatctgcaccaccggcaagctgcc<br>cgtgccttggcctaccctcgtgaccacactgacctacggcgt<br>gcagtgcttcgccagataccccgaccacatgaagcagca<br>cgatttcttcaagagcgccatgcccgagggctacgtgcagg<br>aacggaccatcttcttcaaggacgacggcaactacaagac<br>aagagccgaagtgaagttcgagggcgacaccctcgtgaa<br>ccggatcgagctgaagggcatcgacttcaaagaggatgg<br>caacatcctgggccacaagctggagtacaactacaacag<br>ccacaaggtgtacatcaccgccgacaagcagaaaaacg<br>gcatcaaagtgaacttcaagacccggcacaacatcgagg<br>acggcagcgtgcagctggccgaccactaccagcagaac<br>acccccatcggagatggccccgtgctgctgcccgacaacc<br>actacctgagcacacaaagcgccctgagcaaggacccc<br>aacgagaagcgggaccacatggtgctgctggaatttgtga<br>ccgccgctggcatcaccctgggcatggacgagctgtacaa<br>gtgataatactaaatcgcactgtcggcgtccccccctaacgt<br>tactggccgaagccgcttggaataaggccggtgtgcgtttgt<br>ctatatgttattttccaccatattgccgtcttttggcaatgtgagg<br>gcccggaaacctggccctgtcttcttgacgagcattcctagg<br>ggtctttcccctctcgccaaaggaatgcaaggtctgttgaat<br>gtcgtgaaggaagcagttcctctggaagcttcttgaagaca<br>aacaacgtctgtagcgacccttttgcaggcagcggaacccc |

| | | | |
|---|---|---|---|
| | | | ccacctggcgacaggtgcctctgcggccaaaagccacgt<br>gtataagatacacctgcaaaggcggcacaaccccagtgc<br>cacgttgtgagttggatagttgtggaaagagtcaaatggctc<br>tcctcaagcgtattcaacaaggggctgaaggatgcccaga<br>aggtaccccattgtatgggatctgatctggggcctcggtgca<br>catgctttacatgtgtttagtcgaggttaaaaaacgtctaggc<br>cccccgaaccacggggacgtggttttcctttgaaaaacacg<br>atgataatatggccacaaccatggtgagcaagggcgagg<br>aggataacatggccatcatcaaggagttcatgcgcttcaag<br>gtgcacatggagggctccgtgaacggccacgagttcgag<br>atcgagggcgagggcgagggccgcccctacgagggcac<br>ccagaccgccaagctgaaggtgaccaagggcggccccc<br>tgcccttcgccctgggacatcctgtcccctcagttcatgtacg<br>gctccaaggcctacgtgaagcaccccgccgacatccccg<br>actacttgaagctgtccttccccgagggcttcaagtgggagc<br>gcgtgatgaacttcgaggacggcggcgtggtgaccgtgac<br>ccaggactcctccctgcaggacggcgagttcatctacaag<br>gtgaagctgcgcggcaccaacttcccctccgacggccccg<br>taatgcagaagaagaccatgggctgggaggcctcctccg<br>agcggatgtaccccgaggacggcgccctgaagggcgag<br>atcaagcagaggctgaagctgaaggacggcggccacta<br>cgacgctgaggtcaagaccacctacaaggccaagaagc<br>ccgtgcagctgcccggcgcctacaacgtcaacatcaagtt<br>ggacatcacctcccacaacgaggactacaccatcgtgga<br>acagtacgaacgcgccgagggccgccactccaccggcg<br>gcatggatgagctgtacaagtgaaggatccctcgaggggc<br>ccaagcttacgcgtgcatgcgacgtcatagctctctccctat<br>agtgagtcgtattataagctagcttgggatctttgtgaaggaa<br>ccttacttctgtggtgtgacataattggacaaactacctacag<br>agatttaaagctctaaggtaaatataaaatttttaagtgtataa<br>tgtgttaaactagctgcatatgcttgctgcttgagagttttgctta<br>ctgagtatgatttatgaaaatattatacacaggagctagtgat<br>tataattgtttgtgtattttagattcacagtcccaaggctcatttc<br>aggcccctcagtcctcacagtctgttcatgatcataatcagc<br>cataccacatttgtagaggttttacttgctttaaaaaacctccc<br>acacctccccctgaacctgaaacataaaatgaatgcaattg<br>ttgttgttcgtggctgaatcaacctctggattacaaaatttgtga<br>aagattgactggtattcttaactatgttgctccttttacgctatgt<br>ggatacgctgctttaatgcctttgtatcatgctattgcttcccgt<br>atggctttcattttctcctccttgtataaatcctggttgctgtctctt<br>atgaggagttgtggcccgttgtcaggcaacgtggcgtggtgt<br>gcactgtgtttgctgacgcaacccccactggttggggcattg<br>ccaccacctgtcagctcctttccgggactttcgctttccccctc<br>cctattgccacggcggaactcatcgccgcctgccttgcccg<br>ctgctggacaggggctcggctgttgggcactgacaattccg<br>tggtgttgtcggggaaatcatcgtcctttccttggctgctcgcct<br>gtgttgccacctggattctgcgcgggacgtccttctgctacgt<br>cccttcggccctcaatccagcggaccttccttcccgcggcct<br>gctgccggctctgcggcctcttccgcgtcttcgccttcgccct<br>cagacgagtcggatctccctttgggccgcctccccgccaga<br>gacaattaacttcgcgggtggcatccctgtgacccctcccc<br>agtgcctctcctggccctggaagttgccactccagtgcccac<br>cagccttgtcctaataaaattaagttgcatcattttgtctgacta<br>ggtgtccttctataatattatggggtggaggggggtggtatgg<br>agcaaggggcaagttgggaagacaacctgtagggcctgc<br>ggggtctattgggaaccaagctggagtgcagtggcacaat<br>cttggctcactgcaatctccgcctcctgggttcaagcgattct<br>cctgcctcagcctcccgagttgttgggattccaggcatgcat<br>gaccaggctcagctaatttttgtttttttggtagagacggggttt<br>caccatattggccaggctggtctccaactcctaatctcaggt<br>gatctacccaccttggcctcccaaattgctgggattacaggc<br>gtgaaccactgctcccttccctgtccttgaagttctcagatcct<br>gcattaatgaatcggccaacgcgcggggagaggcggtttg<br>cgtattggctggcgtaatagcgaagaggcccgcaccgatc<br>gcccttcccaacagttgcgcagcctgaatggcgaatggga |

| | | | |
|---|---|---|---|
| | | | cgcgccctgtagcggcgcattaagcgcggcgggtgtggtg<br>gttacgcgcagcgtgaccgctacacttgccagcgccctag<br>cgcccgctcctttcgctttcttcccttcctttctcgccacgttcgc<br>cggctttccccgtcaagctctaaatcgggggctccctttagg<br>gttccgatttagtgctttacggcacctcgaccccaaaaaactt<br>gattagggtgatggttcacgtagtgggccatcgccctgatag<br>acggtttttcgccctttgacgttggagtccacgttcttaatagtg<br>gactcttgttccaaactggaacaacactcaaccctatctcgg<br>tctattcttttgatttataagggattttgccgatttcggcctattggt<br>taaaaaatgagctgatttaacaaaaatttaacgcgaattttta<br>acaaaatattaacgcttacaatttaggtggcacttttcgggga<br>aatgtgcgcggaacccctatttgtttatttttctaaatacattca<br>aatatgtatccgctcatgccaggtcttggactggtgagaacg<br>gcttgctcggcagcttcgatgtgtgctggagggagaataaa<br>ggtctaagatgtgcgatagagggaagtcgcattgaattatgt<br>gctgtgtagggatcgctggtatcaaatatgtgtgcccacccc<br>tggcatgagacaataaccctgataaatgcttcaataatattg<br>aaaaaggaagagtatgagtattcaacatttccgtgtcgccct<br>tattcccttttttgcggcattttgccttcctgtttttgctcacccaga<br>aacgctggtgaaagtaaaagatgctgaagatcagttgggt<br>gcacgagtgggttacatcgaactggatctcaacagcggta<br>agatccttgagagttttcgccccgaagaacgttttccaatgat<br>gagcacttttaaagttctgctatgtggcgcggtattatcccgta<br>ttgacgccgggcaagagcaactcggtcgccgcatacacta<br>ttctcagaatgacttggttgagtactcaccagtcacagaaaa<br>gcatcttacggatggcatgacagtaagagaattatgcagtg<br>ctgccataaccatgagtgataacactgcggccaacttacttc<br>tgacaacgatcggaggaccgaaggagctaaccgcttttttg<br>cacaacatgggggatcatgtaactcgccttgatcgttggga<br>accggagctgaatgaagccataccaaacgacgagcgtg<br>acaccacgatgcctgtagcaatggcaacaacgttgcgcaa<br>actattaactggcgaactacttactctagcttcccggcaaca<br>attaatagactggatggaggcggataaagttgcaggacca<br>cttctgcgctcggcccttccggctggctggtttattgctgataa<br>atctggagccggtgagcgtgggtctcgcggtatcattgcagt<br>actggggccagatggtaagccctcccgtatcgtagttatcta<br>cacgacggggagtcaggcaactatggatgaacgaaatag<br>acagatcgctgagataggtgcctcactgattaagcattggta<br>actgtcagaccaagtttactcatatatactttagattgatttaaa<br>acttcatttttaatttaaaaggatctaggtgaagatcctttttgat<br>aatctcatgccataacttcgtataatgtatgctatacgaagtta<br>tggcatgaccaaaatcccttaacgtgagttttcgttccactga<br>gcgtcagaccccgtagaaaagatcaaaggatcttc |
| Full Plasmid | IVT backbone plasmid | Plasmids with this architecture were used to amplify linear DNA templates for IVT reactions. This plasmid was used for creating linear templates for ACC/ACC 2-ORF. | GACATTGATTATTGACTAGTTATTAATAG<br>TAATCAATTACGGGGTCATTAGTTCATAG<br>CCCATATATGGAGTTCCGCGTTACATAAC<br>TTACGGTAAATGGCCCGCCTGGCTGACC<br>GCCCAACGACCCCCGCCCATTGACGTCA<br>ATAATGACGTATGTTCCCATAGTAACGCC<br>AATAGGGACTTTCCATTGACGTCAATGG<br>GTGGAGTATTTACGGTAAACTGCCCACT<br>TGGCAGTACATCAAGTGTATCATATGCCA<br>AGTACGCCCCCTATTGACGTCAATGACG<br>GTAAATGGCCCGCCTGGCATTATGCCCA<br>GTACATGACCTTATGGGACTTTCCTACTT<br>GGCAGTACATCTACGTATTAGTCATCGCT<br>ATTACCATGGTGATGCGGTTTTGGCAGT<br>ACATCAATGGGCGTGGATAGCGGTTTGA<br>CTCACGGGGATTTCCAAGTCTCCACCCC<br>ATTGACGTCAATGGGAGTTTGTTTTGGCA<br>CCAAAATCAACGGGACTTTCCAAAATGTC<br>GTAACAACTCCGCCCCATTGACGCAAAT<br>GGGCGGTAGGCGTGTACGGTGGGAGGT |

| | | | |
|---|---|---|---|
| | | | CTATATAAGCAGAGCTggtttagtgaaccgtcag atccgctagagatccgcggccgctaatacgactcactatag ggtcagatcgcctggagacgccatccacgctgttttgacctc catagaagacaccgggaccgatccagcctccggactcta gcctaggcttttgcaaaaagctatttaggtgacactatagaa ggtacgcctgcaggtaccgagctcggatccagtagtttaaa ccccttcaccatggtgTCCAAGGGCGAGGAACT GTTCACCGGCGTGGTGCCCATCCTGGTG GAACTGGACGGCGACGTGAACGGCCAC AAGTTCAGCGTGAGAGGCGAGGGCGAA GGCGACGCCACAAACGGAAAGCTGACC CTGAAGTTCATCTGCACCACCGGCAAGC TGCCCGTGCCTTGGCCTACCCTCGTGAC CACACTGACCTACGGCGTGCAGTGCTTC AGCAGATACCCCGACCACATCAAGAGAC ACGATTTCTTCAAGAGCGCCCTGCCCGA GGGCTACGTGCAGGAACGGACCATCAG CTTCAAGGACGACGGCACCTACAAGACA AGAGCCGAAGTGAAGTTCGAGGGCGAC ACCCTCGTGAACCGGATCGAGCTGAAGG GCATCGACTTCAAAGAGGACGGCAACAT CCTGGGCCACAAGCTGGAGTACAACTTC AACAGCCACAACGTGTACATCACCGCCG ACAAGCAGAAAAACGGCATCAAAGCCAA CTTCAAGATCCGGCACAACGTGGAGGAC GGCAGCGTGCAGCTGGCCGACCACTAC CAGCAGAACACCCCCATCGGAGACGGC CCCGTGCTGCTGCCCGACAACCACTACC TGAGCACACAAAGCAAGCTGAGCAAGGA CCCCAACGAGAAGCGGGACCACGCCGT GCTGCTGGAATTTGTGACCGCCGCTGGC ATCACCCACggcaaggacgagctgtacaagtgacc cgggttaccatggtgtccaagggcgaggaactgttcaccg gcgtggtgcccatcctggtggaactggatggcgacgtgaa cggccacaagttctctgtgcggggagagggcgaaggcga cgccacaaatggcaagctgaccctgaagttcatctgcacc accggcaagctgcccgtgccttggcctaccctcgtgaccac actgtctcacggcgtgcagtgcttcgccagataccccgacc acatgaagcagcacgatttcttcaagagcgccatgcccga gggctacgtgcaggaacggaccatcttcttcaaggacgac ggcacctacaagaccagagccgaagtgaagttcgaggg cgacaccctcgtgaaccggatcgagctgaagggcgtgga cttcaaagaggacggcaacatcctgggccacaagctgga gtacaacttcaacagccacaacatctacatcatggccgtga agcagaaaaacggcatcaaagtgaacttcaagatccggc acaacgtggaagatggcagcgtgcagctggccgaccact accagcagaacacccccatcggagatggccccgtgctgc tgcctgatagccactacctgagcacccagagcaagctgag caaggaccccaacgagaagcgggaccacatggtgctgct ggaatttcggaccgccgctggcatcaccctgggcatggatg agctgtacaagtgataatacCTCGAGCTGGTACT GCATGCACGCAATGCTAGCTGCCCCTTT CCCGTCCTGGGTACCCCGAGTCTCCCCC GACCTCGGGTCCCAGGTATGCTCCCACC TCCACCTGCCCCACTCACCACCTCTGCT AGTTCCAGACACCTCCCAAGCACGCAGC AATGCAGCTCAAAACGCTTAGCCTAGCC ACACCCCCACGGGAAACAGCAGTGATTA ACCTTTAGCAATAAACGAAAGTTTAACTA AGCTATACTAACCCCAGGGTTGGTCAAT TTCGTGCCAGCCACACCCTGGAGCTAGC AAAAAAAAAAAAAAAAAAAAAAAAAAAAA AGCATATGACTAAAAAAAAAAAAAAAAAA AAAAAAAAAAAAAAAAAAAAAAAAAAAAA AAAAAAAAAAAAAAAAAAAAAAAGAGACC |

| | | | |
|---|---|---|---|
| | | | aataaaagatctttattttcattagatctgtgtgttggttttttgtgt<br>gGGCTGGATGATCCTCCAGCGCGGGGAT<br>CTCATGCTGGAGTTCTTCGCCCACCCCA<br>ACTTGTTTATTGCAGCTTATAATGGTTAC<br>AAATAAAGCAATAGCATCACAAATTTCAC<br>AAATAAAGCATTTTTTTCACTGCATTCTA<br>GTTGTGGTTTGTCCAAACTCATCAATGTA<br>TCTTATCATGTCTGTATACCGTCGACCTC<br>TAGCTAGAGCTTGGCGTAATCATGGTCA<br>TAGCTGTTTCCTGTGTGAAATTGTTATCC<br>GCTCACAATTCCACACAACATACGAGCC<br>GGAAGCATAAAGTGTAAAGCCTGGGGTG<br>CCTAATGAGTGAGCTAACTCACATTAATT<br>GCGTTGCGCTCACTGCCCGCTTTCCAGT<br>CGGGAAACCTGTCGTGCCAGCTGCATTA<br>ATGAATCGGCCAACGCGCGGGGAGAGG<br>CGGTTTGCGTATTGGGCGCTCTTCCGCT<br>TCCTCGCTCACTGACTCGCTGCGCTCGG<br>TCGTTCGGCTGCGGCGAGCGGTATCAG<br>CTCACTCAAAGGCGGTAATACGGTTATC<br>CACAGAATCAGGGGATAACGCAGGAAAG<br>AACATGTGAGCAAAAGGCCAGCAAAAGG<br>CCAGGAACCGTAAAAAGGCCGCGTTGCT<br>GGCGTTTTTCCATAGGCTCCGCCCCCCT<br>GACGAGCATCACAAAAATCGACGCTCAA<br>GTCAGAGGTGGCGAAACCCGACAGGAC<br>TATAAAGATACCAGGCGTTTCCCCCTGG<br>AAGCTCCCTCGTGCGCTCTCCTGTTCCG<br>ACCCTGCCGCTTACCGGATACCTGTCCG<br>CCTTTCTCCCTTCGGGAAGCGTGGCGCT<br>TTCTCATAGCTCACGCTGTAGGTATCTCA<br>GTTCGGTGTAGGTCGTTCGCTCCAAGCT<br>GGGCTGTGTGCACGAACCCCCCGTTCA<br>GCCCGACCGCTGCGCCTTATCCGGTAAC<br>TATCGTCTTGAGTCCAACCCGGTAAGAC<br>ACGACTTATCGCCACTGGCAGCAGCCAC<br>TGGTAACAGGATTAGCAGAGCGAGGTAT<br>GTAGGCGGTGCTACAGAGTTCTTGAAGT<br>GGTGGCCTAACTACGGCTACACTAGAAG<br>aACAGTATTTGGTATCTGCGCTCTGCTGA<br>AGCCAGTTACCTTCGGAAAAAGAGTTGG<br>TAGCTCTTGATCCGGCAAACAAACCACC<br>GCTGGTAGCGGTGGTTTTTTTGTTTGCAA<br>GCAGCAGATTACGCGCAGAAAAAAAGGA<br>TCTCAAGAAGATCCTTTGATCTTTTCTAC<br>GGGGTCTGACGCTCAGTGGAACGAAAAC<br>TCACGTTAAGGGATTTTGGTCATGAGATT<br>ATCAAAAAGGATCTTCACCTAGATCCTTT<br>TAAATTAAAAATGAAGTTTTAAATCAATCT<br>AAAGTATATATGAGTAAACTTGGTCTGAC<br>AGTTACCAATGCTTAATCAGTGAGGCAC<br>CTATCTCAGCGATCTGTCTATTTCGTTCA<br>TCCATAGTTGCCTGACTCCCCGTCGTGT<br>AGATAACTACGATACGGGAGGGCTTACC<br>ATCTGGCCCCAGTGCTGCAATGATACCG<br>CGAGACCCACGCTCACCGGCTCCAGATT<br>TATCAGCAATAAACCAGCCAGCCGGAAG<br>GGCCGAGCGCAGAAGTGGTCCTGCAAC<br>TTTATCCGCCTCCATCCAGTCTATTAATT<br>GTTGCCGGGAAGCTAGAGTAAGTAGTTC<br>GCCAGTTAATAGTTTGCGCAACGTTGTT<br>GCCATTGCTACAGGCATCGTGGTGTCAC<br>GCTCGTCGTTTGGTATGGCTTCATTCAG<br>CTCCGGTTCCCAACGATCAAGGCGAGTT<br>ACATGATCCCCCATGTTGTGCAAAAAAG |

| | | | |
|---|---|---|---|
| | | | CGGTTAGCTCCTTCGGTCCTCCGATCGT<br>TGTCAGAAGTAAGTTGGCCGCAGTGTTA<br>TCACTCATGGTTATGGCAGCACTGCATA<br>ATTCTCTTACTGTCATGCCATCCGTAAGA<br>TGCTTTTCTGTGACTGGTGAGTACTCAAC<br>CAAGTCATTCTGAGAATAGTGTATGCGG<br>CGACCGAGTTGCTCTTGCCCGGCGTCAA<br>TACGGGATAATACCGCGCCACATAGCAG<br>AACTTTAAAAGTGCTCATCATTGGAAAAC<br>GTTCTTCGGGGCGAAAACTCTCAAGGAT<br>CTTACCGCTGTTGAGATCCAGTTCGATG<br>TAACCCACTCGTGCACCCAACTGATCTT<br>CAGCATCTTTTACTTTCACCAGCGTTTCT<br>GGGTGAGCAAAAACAGGAAGGCAAAATG<br>CCGCAAAAAAGGGAATAAGGGCGACAC<br>GGAAATGTTGAATACTCATACTCTTCCTT<br>TTTCAATATTATTGAAGCATTTATCAGGG<br>TTATTGTCTCATGAGCGGATACATATTTG<br>AATGTATTTAGAAAAATAAACAAATAGGG<br>GTTCCGCGCACATTTCCCCGAAAAGTGC<br>CACCTGACGTCGACGGATCGGGAGATCT<br>CCCGATCCCCTATGGTGCACTCTCAGTA<br>CAATCTGCTCTGATGCCGCATAGTTAAG<br>CCAGTATCTGCTCCCTGCTTGTGTGTTG<br>GAGGTCGCTGAGTAGTGCGCGAGCAAA<br>ATTTAAGCTACAACAAGGCAAGGCTTGA<br>CCGACAATTGCATGAAGAATCTGCTTAG<br>GGTTAGGCGTTTTGCGCTGCTTCGTACG<br>GGCCAGATATACGCGTT |
